# Inhibition of 2-Hydroxyglutrate Elicits Metabolic-reprograming and Mutant IDH1 Glioma Immunity

**DOI:** 10.1101/2020.05.11.086371

**Authors:** Padma Kadiyala, Stephen V. Carney, Jessica C. Gauss, Maria B. Garcia-Fabiani, Felipe J. Núñez, Fernando M. Nunez, Mahmoud S. Alghamri, Yayuan Liu, Minzhi Yu, Dan Li, Marta B. Edwards, James J. Moon, Anna Schwendeman, Pedro R. Lowenstein, Maria G. Castro

**Affiliations:** Department of Neurosurgery, University of Michigan Medical School, Ann Arbor, MI 48109, USA; Department of Cell and Developmental Biology, University of Michigan Medical School, Ann Arbor, MI 48109, USA; Department of Pharmaceutical Sciences, University of Michigan, Ann Arbor, MI 48109, USA; Biointerfaces Institute, University of Michigan Medical School, Ann Arbor, MI 48109, USA; Department of Biomedical Engineering, University of Michigan, Ann Arbor, MI 48109, USA

**Keywords:** Mutant IDH1, genetically engineered glioma models, IDH1-R132H Inhibitors, Immunotherapy.

## Abstract

Mutant isocitrate-dehydrogenase-1 (IDH1-R132H; mIDH1) is a hallmark of adult gliomas. Lower grade mIDH1 gliomas are classified into two molecular subgroups: (i) 1p/19q co-deletion/TERT-promoter mutations or (ii) inactivating mutations in α-thalassemia/mental retardation syndrome X-linked (*ATRX*) and *TP53.* This work, relates to the gliomas’ subtype harboring mIDH1, *TP53* and *ATRX* inactivation. IDH1-R132H is a gain-of-function mutation that converts α-ketoglutarate into 2-hydroxyglutarate (D-2HG). The role of D-2HG within the tumor microenvironment of mIDH1/mATRX/mTP53 gliomas remains unexplored. Inhibition of 2HG, when used as monotherapy or in combination with radiation and temozolomide (IR/TMZ), led to increased median survival (MS) of mIDH1 glioma bearing mice. Also, 2HG inhibition elicited anti-mIDH1 glioma immunological memory. In response to 2HG inhibition, PD-L1 expression levels on mIDH1-glioma cells increased to similar levels as observed in wild-type-IDH1 gliomas. Thus, we combined 2HG inhibition/IR/TMZ with anti-PDL1 immune checkpoint-blockade and observed complete tumor regression in 60% of mIDH1 glioma bearing mice. This combination strategy reduced T-cell exhaustion and favored the generation of memory CD8^+^T-cells. Our findings demonstrate that metabolic reprogramming elicits anti-mIDH1 glioma immunity, leading to increased MS and immunological memory. Our preclinical data supports the testing of IDH-R132H inhibitors in combination with IR/TMZ and anti-PDL1 as targeted therapy for mIDH1/mATRX/mTP53 glioma patients.

**Brief Summary:** Inhibition of 2-Hydroxyglutrate in mutant-IDH1 glioma in the genetic context of ATRX and TP53 inactivation elicits metabolic-reprograming and anti-glioma immunity.

## Introduction

Gliomas are highly infiltrative brain tumors accounting for 32% of all primary central nervous system malignancies (1). With advances in molecular biology and sequencing technologies, a distinct profile of genetic alterations for gliomas has emerged (1–3). A gain-of-function mutation in the gene encoding isocitrate dehydrogenase 1 (IDH1) mutation has been reported in ∼46% of all adult gliomas (2) and ∼80% of low grade gliomas (3, 4). This mutation results in the replacement of arginine (R) for histidine (H) at amino acid residue 132 (R132H) (5, 6). In gliomas, the IDH1-R132H mutation co-occurs with the following genetic alterations: i) oligodendroglioma-1p/19q co-deletion, *TERT* promoter mutations, or ii) astrocytoma-inactivation of tumor suppressor protein 53 *(TP53)* gene and loss of function mutations in alpha thalassemia/mental retardation syndrome X-linked gene *(ATRX)*(2, 4, 7).

The IDH1-R132H neomorphic mutation (mIDH1) confers a gain-of-function catalytic activity, prompting the NADPH-dependent reduction of alpha ketoglutarate (α-KG) to the oncometabolite D-2-hydroxyglutarate (2-HG) (5, 8, 9). The accumulation of 2-HG acts antagonistically to α-KG, competitively inhibiting α-KG-dependent dioxygenases, including the ten-eleven translocation (TET) methylcytosine dioxygenases and Jumonji C (JmjC) domain-containing histone demethylases (5, 9–11). This leads to a DNA and histone H3 hypermethylation phenotype resulting in an epigenetic reprogramming of the glioma cell transcriptome (7, 12, 13).

Ongoing research has demonstrated the benefits of targeting IDH1-R132H in gliomas with small molecule inhibitors (14). The compound AGI-5198 is an allosteric, competitive inhibitor that is selective for the IDH1-R132H enzyme, inhibiting the synthesis of 2-HG in mouse and human glioma cells (7, 15, 16). Our laboratory has generated a genetically engineered mouse model of glioma expressing IDH1-R132H (mIDH1) concomitantly with loss of ATRX and TP53 to study IDH1-R132H within the scope of the genetic lesions encountered in human astrocytomas (2, 7). Using this model, we demonstrated that AGI-5198 treatment decreased 2-HG levels, induced radiosensitivity and decreased proliferation of mIDH1 glioma neurospheres *in vitro* (7). AGI-5198 has also been shown to inhibit 2-HG production *in vivo* and impair tumor growth in an anaplastic oligodendroglioma patient derived xenograft model expressing 1p/19q co-deletion and IDH1-R132H (15). AGI-5198 has been structurally optimized to AG-120 for clinical evaluation for the treatment of cholangiocarcinoma, chondrosarcoma, and acute myeloid leukemia (AML) (16). AG-120 is currently in late-stage clinical development for AML (NCT03173248) and cholangiocarcinoma (NCT02989857) (16). Recently, AG-120 has been shown to have a favorable safety profile for non-enhancing IDH1-R132H gliomas (17).

Herein, we used our mouse model of glioma expressing IDH1-R132H and loss of ATRX and TP53 (mIDH1) (7) to elucidate the role of D-2HG in the glioma immune microenvironment. Our findings demonstrate that treating mIDH1 glioma bearing mice with IDH1-R132H inhibitor alone or in combination with radiation (IR) and temozolomide (TMZ), which is standard of care (SOC) for mIDH1 glioma patients, significantly prolonged the median survival (MS) of mIDH1 glioma bearing mice and elicited anti-glioma immunity. IDH1-R132H inhibition used in combination with SOC increased the frequency of tumor-specific cytotoxic CD8^+^ T cells and interferon-γ (IFN-γ) release within the TME. Strikingly, long-term survivors from IDH1-R132H inhibition/SOC treatment group remained tumor-free after rechallenging them with mIDH1 glioma in the contralateral hemisphere, indicating the development of anti-mIDH1 glioma immunological memory. This is a critical factor in determining the success of immune-therapeutic approaches in gliomas. A robust anti-tumor T cell response and the presence of anti-glioma immunological memory are required to eradicate any remnant tumor cells post-surgery and prevent recurrence.

We observed that genetically engineered mIDH1 mouse gliomas, resembling human mutant IDH1 astrocytoma, exhibit lower levels of PD-L1 expression on the tumor cells’ surface. In response to 2HG inhibition, PD-L1 expression levels on mIDH1 glioma cells significantly increased to those observed in wild type IDH gliomas. Numerous preclinical solid tumor models have demonstrated that the immune checkpoint blockade of PD-1/PD-L1 interaction prevents T cell exhaustion, resulting in enhanced anti-tumor immune activity and improved MS (18). We previously demonstrated that PD-L1 checkpoint blockade as monotherapy elicited a small increase in MS in mice bearing syngeneic glioma, with only a few long term survivors (19). However, immune-checkpoint blockade used as monotherapy has failed in Phase III clinical trials to improve overall survival of patients with glioma (20). To date, the efficacy of PD-1/PD-L1 immune checkpoint blockade therapy has not been unexplored in glioma models harboring IDH1-R132H in the context of ATRX and TP53 loss. In this study we demonstrate that co-administering anti-PD-L1 immune checkpoint blockade with IDH1-R132H inhibition and SOC significantly enhances the overall survival of mIDH1 glioma bearing mice. Furthermore, this treatment strategy reduces T cell exhaustion and strongly promotes the generation of memory CD8^+^ T cells, leading to immunological memory. Collectively, our findings demonstrate that upon metabolic reprogramming it is possible to elicit anti-glioma immunity, leading to increased MS and anti-tumor immunological memory. Our data supports the clinical testing of IDH1-R132H inhibitors in combination with SOC and anti-PD-L1 immune-checkpoint blockade to treat glioma patients expressing IDH1-R132H in the context of *TP53, ATRX* inactivating mutations.

## Results

### Inhibition of 2HG Production Sensitizes mIDH1 Glioma to Radiotherapy and Induces Immunogenic Cell Death

Previous work from our laboratory demonstrated that gliomas expressing IDH1-R132H (mIDH1) concomitantly with loss of ATRX and TP53 are radioresistant (7). In this study, we sought to determine if inhibiting D-2HG production would sensitize mIDH1 glioma to radiation (IR) treatment and induce immunogenic cell death (ICD). We performed a clonogenic assay on mIDH1 and wtIDH mouse neurospheres (mouse-NS) (Figure 1A) and human cells (human-GC), by treating them with the mIDH1 inhibitor AGI-5198 and IR. Compared to mIDH1 mouse-NS subjected to IR alone, we observed a significant reduction in clonogenic survival of mIDH1 mouse-NS treated with AGI-5198 and IR (*P* ≤ 0.0001), suggesting that AGI-5198 treatment radiosensitizes mIDH1 NS (Figure 1B). On the other hand, we observed a dose dependent decrease in clonogenic survival of wtIDH mouse-NS in response to IR, irrespective of AGI-5198 treatment (Figure 1B). Notably, treating radioresistant human mIDH1 cells (MGG119) which endogenously express IDH1-R132H, *ATRX* and *TP53* inactivating mutations with AGI-5198 rendered them radiosensitive (*P* ≤ 0.0001; Figure 1C). AGI-5198 treatment of radiosensitivity human wtIDH (SJGBM2) cells which endogenously express *ATRX* and *TP53* inactivating mutations did not alter their response to radiotherapy (Figure 1C).

**Figure 1:**
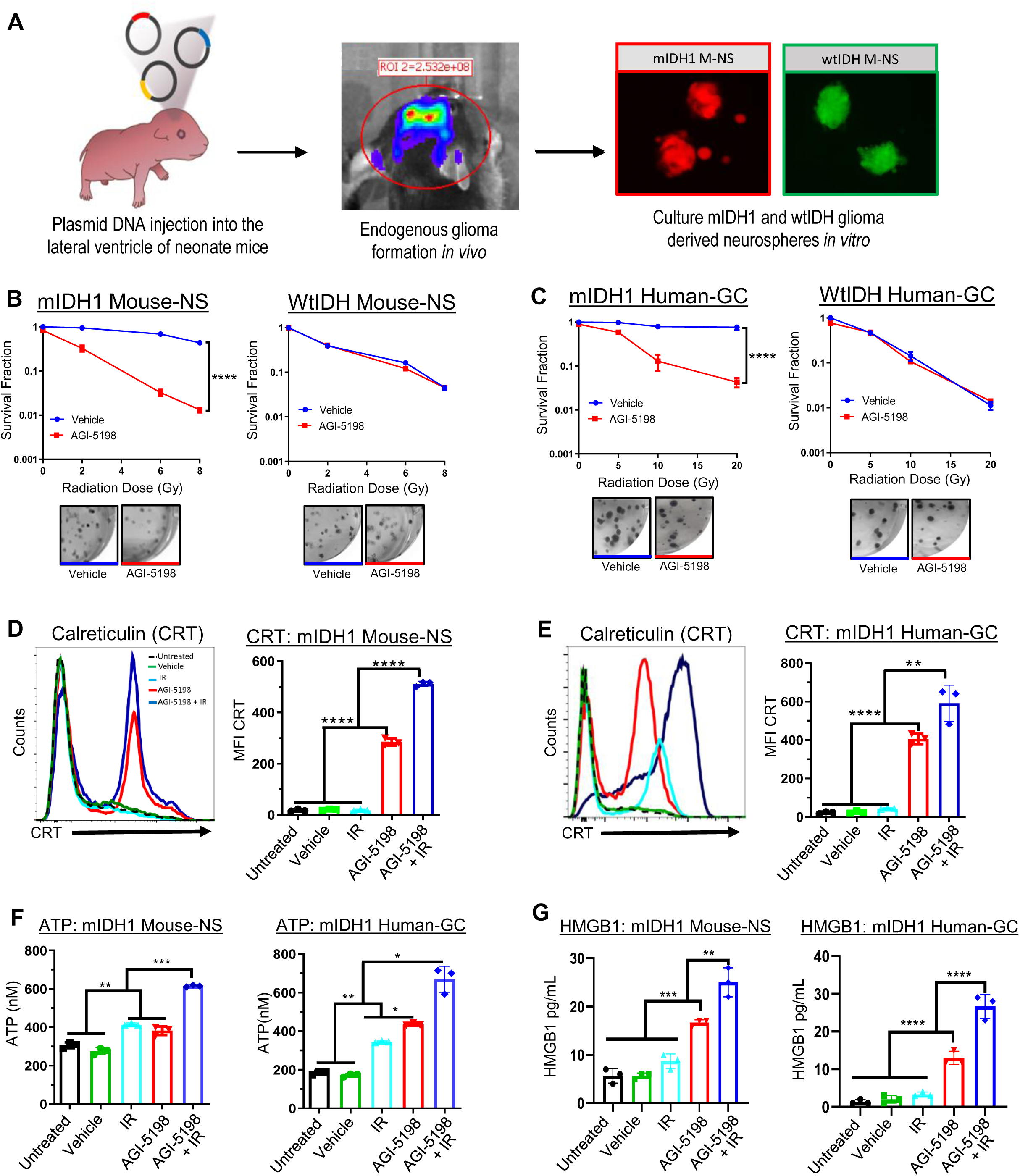
Inhibition of IDH1-R132H in mIDH1 mouse-NS and human glioma cells confers radiosensitivity and promotes release of endogenous DAMPs. (A) Mouse neurosphere (Mouse-NS) cultures were generated from genetically engineered wtIDH1 or mIDH1 glioma mouse models. **(B, C)** Clonogenic assay of wtIDH1 and mIDH1 Mouse-NS and human glioma cells (Human-GC). (**B**) Mouse-NS after treatment with 0 to 8 Gy of ionizing radiation (IR) in the presence of 1.5 μM AGI-5198 (red) or DMSO vehicle (blue). (**C**) Human-GCs SJGBM2 (wtIDH1) and MGG119 (mIDH1) after treatment with 0 to 20 Gy of IR in the presence of 5 μM AGI-5198 (red) or DMSO vehicle (blue). \*\*\*\**P* < 0.0001, Non-Linear Regression. Bars represent mean ± SEM (*n* = 3 technical replicates). (**D-G**) Calreticulin (CRT), ATP, and HMGB1 DAMP molecules’ expression levels within mIDH1 Mouse-NS and Human-GC. Mouse-NS were treated with 3 Gy IR in combination 1.5 μM of AGI-5198 for 72 hrs. Human-GC were treated with 10 Gy IR in combination with 5μM of AGI-5198 for 72 hrs. Quantification of CRT expression on mIDH1 Mouse-NS and Human-GC after treatment is shown in D and E, respectively. Representative histograms display CRT marker’s expression levels (black = Untreated, green = DMSO Vehicle, light blue=IR, red=AGI-5198, and dark blue = AGI5198 + IR). \*\*\*\**P* < 0.0001, \*\**P* < 0.001, one-way ANOVA test. Bars represent mean ± SEM (*n* = 3 technical replicates). (**F**) Quantification of ATP release in the supernatant of mIDH1 Mouse-NS and Human-GC. \**P* < 0.01; \*\**P* < 0.01; \*\*\**P* < 0.001, one-way ANOVA test. Bars represent mean ± SEM (*n* = 3 technical replicates). (**G**) Quantification of HMGB1 release in the supernatant of mIDH1 Mouse-NS and Human-GC. \*\**P* < 0.01; \*\*\**P* < 0.001; \*\*\*\**P* < 0.0001, one-way ANOVA test. Bars represent mean ± SEM (*n* = 3 technical replicates).

Next, we assessed if AGI-5198 therapy in combination with IR induces ICD in mIDH1 mouse-NS and human-GC. We measured the levels of calreticulin (CRT), ATP, and HMGB-1, IL-1α, and IL-6, common damage-associated molecular pattern molecules (DAMPs) expressed by dying tumor cells (21). First, we quantified the levels of CRT expression in mIDH1 mouse-NS and human-GC cells in response AGI-5198 alone, IR alone, or AGI-5198+IR treatments. The mIDH1 mouse-NS treated with AGI-5198 displayed a ∼6-fold (*P* ≤ 0.0001) increase in CRT expression relative to untreated, DMSO control, and IR alone groups (Figure 1D). This response was further increased by ∼1.5-fold (*P* ≤ 0.0001) with AGI-5198+IR treatment (Figure 1D). Notably, in mIDH1 human-GC cells, CRT expression was ∼7-fold (*P* ≤ 0.0001) higher for AGI-5198 treatment group compared to untreated, DMSO and IR treatment groups (Figure 1E). When mIDH1 human-GC cells were treated with AGI-5198+IR, there was an additional ∼1.5-fold (*P* ≤ 0.01) increase in the CRT expression on the surface of mIDH1 glioma cells (Figure 1E). We also tested the amount of ATP release in the supernatants of mIDH1 mouse-NS and human-GC in response to if AGI-5198, IR, or AGI-5198+IR treatments. We observed a ∼2-fold (*P* ≤ 0.01) increase in the extracellular release of ATP in the supernatant of mIDH1 mouse-NS and human-GCs treated with AGI-5198 compared to untreated or DMSO vehicle treatment groups (Figure 1F). We observed an additional ∼1.5-fold (*P* ≤ 0.001) increase in extra cellular ATP concentration in the supernatant of mIDH1 mouse-NS and human-GC treated with AGI-5198+IR (Figure 1F). No difference was observed in ATP release between IR and AGI-5198 treatment groups in mIDH1 mouse-NS. However, there was ∼1-fold (*P* ≤ 0.05) increase in ATP release in mIDH1 human-GC treated with AGI-5198 compared to IR treatment group (Figure 1F). Lastly, we tested the release of HMGB1 in the supernatants of mIDH1 mouse-NS and human-GC in response to if AGI-5198, IR, or AGI-5198+IR treatments. We observed a ∼1.5-fold (*P* ≤ 0.001) increase in the extra cellular release of HMGB1 in the supernatant of mIDH1 mouse-NS treated with AGI-5198 compared to untreated, IR or DMSO treatment groups (Figure 1G). We observed an additional ∼1.6-fold (*P* ≤ 0.01) increase in extra cellular HMGB1 release in the supernatant of mIDH1 mouse-NS with AGI-5198+IR (Figure 1G). There was ∼1.5-fold (*P* ≤ 0.0001) increase in the extra cellular release of HMGB1 in the supernatant of mIDH1 human-GC treated with AGI-5198 compared to untreated, DMSO, or IR treatment groups. An additional ∼1.5-fold (*P* ≤ 0.0001) increase in extra cellular HMGB1 release in the supernatant of mIDH1 human-GC treated with AGI-5198+IR was also observed (Figure 1G). We observed similar results for additional DAMPs such as IL-1α (21) and IL-6 (21) in both mIDH1 Mouse NS and Human GC (Supplementary Figure 1). Taken together, these results demonstrate that AGI-5198 therapy in combination with IR induces ICD in mIDH1 mouse-NS and human-GC.

### Pharmacological inhibition of IDH1-R132H Prolongs the Survival of mIDH1 Glioma Bearing Mice

Firstly, we sought to address if tumor-derived D-2HG production can be suppressed by IDH1-R132H inhibitor AGI-5198 *in vivo*. Mice bearing either wtIDH1 or mIDH1 glioma were treated with AGI-5198 or saline as indicated in Figure 2A. Tumors were extracted from mice and processed for UPLC-MS analysis two days following the last treatment dose (Supplementary Figure 2). Total D-2HG concentration was ∼1.5-fold (0.29 μmol/g) higher in mIDH1 tumor tissue compared to untreated normal brain tissue (0.15 μmol/g; Figure 2B). The levels of D-2HG in mIDH1 brain tumor tissue were reduced by ∼2.4 fold (*P* ≤ 0.0001; 0.12 μmol/g) after treatment with AGI-5198 (Figure 2B). Furthermore, D-2HG levels in the mIDH1 brain tumor tissue of mice treated with AGI-5198 were similar to the amount observed in wtIDH1 brain tissue (0.15 μmol/g; Figure 2B). These data demonstrate that AGI-5198 is capable of inhibiting IDH-R132H activity in the brain our mouse glioma model in vivo.

**Figure 2:**
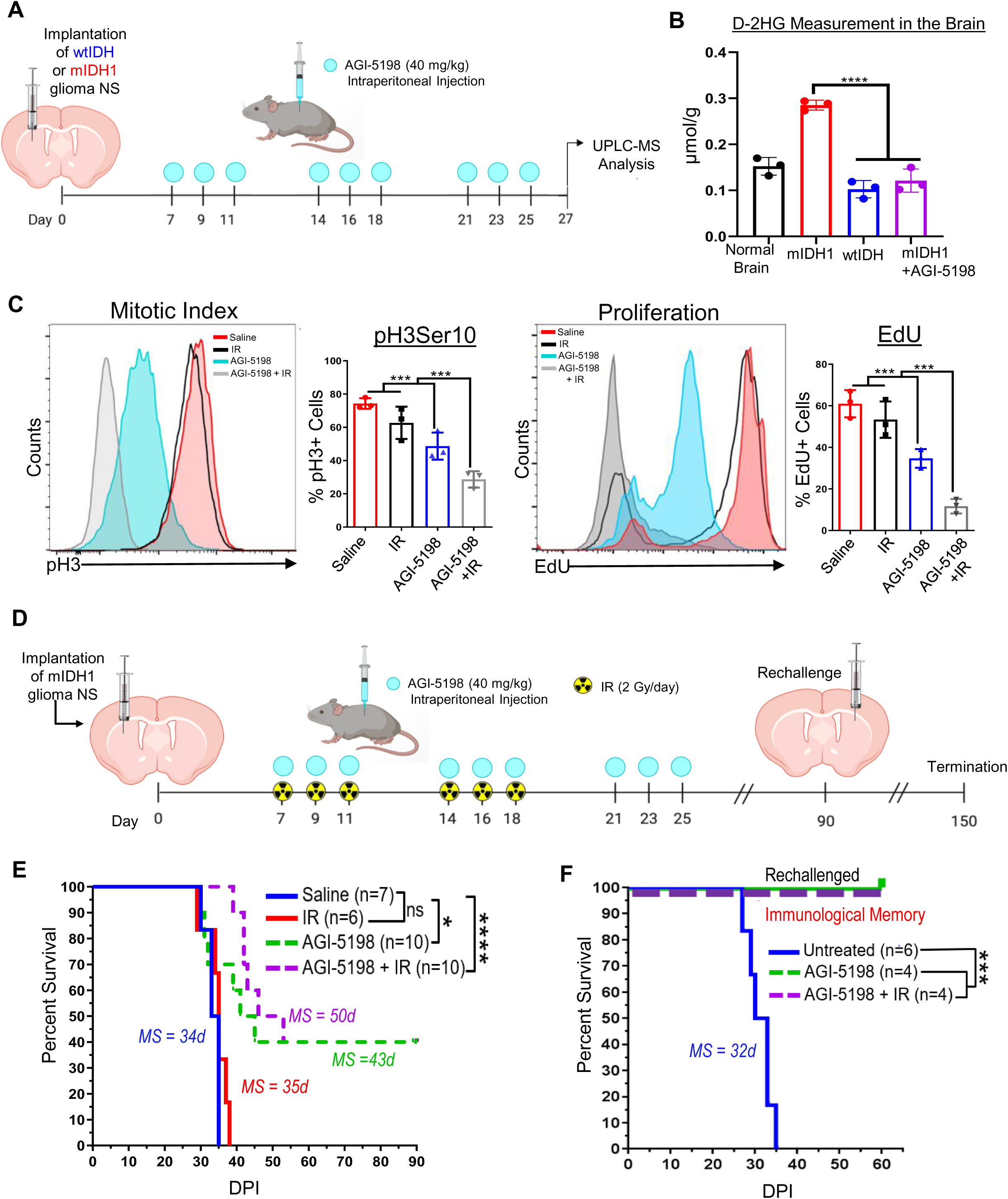
Inhibition of IDH1-R132H decreases the production of 2HG *in vivo*, increases the median survival of mIDH1 glioma bearing mice and elicits anti-mIDH1 immunological memory. (A) Diagram of experimental design to assess D-2HG concentration in the brain tumor microenvironment (TME). At 27 dpi brains were harvested for UPLC-MS analysis of D-2HG in the TME of normal mice (blank; black), wtIDH glioma bearing mice (blue), mIDH1 glioma bearing mice (red), and mIDH1 glioma bearing mice treated with AGI-5198 (purple). (**B**) Quantification of 2HG concentration in normal brain, mIDH1 tumor bearing brain, wtIDH1 tumor bearing brain, and mIDH1 tumor bearing brain plus AGI-5198 treatment. \*\*\*\**P* < 0.0001, two-way ANOVA test. Bars represent mean ± SEM (*n* = 3 biological replicates). (**C**) AGI-5198+IR treatment inhibits mIDH1 glioma growth *in vivo*. Mitotic cell division (mitotic index) in mIDH1 glioma was assessed by the phosphorylation status of Ser10 in Histone 3 (pH3). Proliferation (S phase) of mIDH1 glioma was assessed by measuring the amount of EdU incorporation. Representative histograms display pH3 or EdU positive cells (red = saline, black = IR, light blue = AGI-5198 and grey = AGI-5198 + IR). \*\*\**P* < 0.001, one-way ANOVA test. Bars represent mean ± SEM (*n* = 3 biological replicates). (**D**) Diagram of experimental design to assess survival for AGI-5198 therapy in combination with radiation. (**E)** Kaplan-Meier analysis of saline (*n* = 7), IR (*n* = 6), AGI-5198 (*n* =10), and AGI-5198 + IR (*n* = 10) treated mice. **(F)** Kaplan-Meier plot for rechallenged long-term survivors from (D) AGI-5198 (*n* = 4) or AGI-5198 + IR (*n* = 4), and untreated animals (n=6). Data were analyzed using the log-rank (Mantel-Cox) test. \**P* < 0.05; \*\*\**P* < 0.001; \*\*\*\**P* < 0.0001; MS = median survival; d = days.

Next, we characterized the *in vivo* cell cycle profile of mIDH1 tumor bearing mice treated with AGI-5198, IR, or AGI-5198 in combination with IR. The frequency of mIDH1 cells undergoing mitotic G2/M phase (pH3Ser10^+^) of cell cycle was ∼1.5-fold (*P* ≤ 0.001) lower in the AGI-5198 treatment group compared to the saline control and IR treatment groups (Figure 2C). In mice treated with AGI-5198+IR, we observed ∼1.6 fold (*P* ≤ 0.05; Figure 2C) decrease in the frequency of mIDH1 cells undergoing mitosis compared to AGI-5198 alone treatment group. Actively proliferating mIDH1 cells in S phase (EdU^+^) of cell cycle were ∼2.3-fold (*P* ≤ 0.001) lower in the AGI-5198 treatment group compared to the saline and IR treatment groups (Figure 2C). Mutant IDH1 glioma cell proliferation was further reduced by ∼ 6-fold (*P* ≤ 0.001) in mice treated with AGI-5198+IR (Figure 2C).

Given that D-2HG levels can be reduced *in vivo* (Figure 2B) and having shown that AGI-5198 alone and AGI-5198+IR treatment of mIDH1 glioma cells results in the release of ICD activating DAMPs molecules *in vitro* (Figure 1), we asked whether AGI-5198 alone or AGI-5198+IR will improve the efficacy of mIDH1 glioma bearing mice and elicit anti-mIDH1 glioma immunity. Mutant IDH1 glioma bearing mice were treated with (i) saline, (ii) IR, (iii) AGI-5198, or (iv) AGI-5198+IR at the indicated dose and treatment schedule (Figure 2D). We observed a ∼1.3 fold (*P* ≤ 0.001) increase in median survival (MS) of mice in the AGI-5198 treated group (MS: 43 dpi), and a ∼1.5 fold (*P* ≤ 0.001) increase in MS of mice AGI-5198+IR treated group (MS: 50 dpi), when compared to the control mice in the saline treatment group (MS: 34 dpi) or IR treatment group (MS: 35 dpi). We also evaluated whether the delivery of PEG-400/Ethanol vehicle impacted MS of mIDH1 glioma bearing mice. Treating mIDH1 glioma bearing mice with vehicle control following the same treatment schedule as AGI-5918 (Supplementary Figure 3A) did not confer survival benefit (Supplementary Figure 3B).

Strikingly, we observed that 40% of the mIDH1 bearing mice treated with AGI-5198 or AGI-5198+IR survived long term (> 90 dpi) and remained tumor free (Figure 2E; Table 1). Long-term survivors from AGI-5198 alone or AGI-5198+IR treatment group were rechallenged with mIDH1 mouse-NS in the contralateral hemisphere. These animals remained tumor free without further treatment, whereas control mice implanted with glioma cells succumbed due to tumor burden (MS: 32 days; *P* ≤ 0.0001) (Figure 2F). These results suggest the development of immunological memory in mIDH1 glioma rechallenged animals that were previously treated with AGI-5198 or AGI-5198+IR.

**Table 1:** Log-rank (Mantel-Cox) Test Kaplan Meir Survival Analysis for AGI-5198 treatment in combination with IR

| Group | P value |
| --- | --- |
| Saline vs. IR | 0.2538 |
| Saline vs. AGI-5198 | 0.0406 |
| Saline vs. AGI5198 + IR | <0.0001 |
| AGI-5198 vs. AGI5198 + IR | 0.5818 |

We also assessed the therapeutic efficacy of AGI-5198 treatment in a wtIDH glioma model. Mice bearing wtIDH glioma were treated with (i) saline, (ii) IR, (iii) AGI-5918, and (iv) AGI5198+IR at the indicated doses and treatment schedule (Supplementary Figure 4A). We observed a ∼2.2 fold (*P* ≤ 0.01) increase in MS of wtIDH glioma mice in the IR (MS: 54 dpi), as well as AGI-5198+IR in treatment groups (MS: 51 dpi), when compared to the control mice in the saline (MS: 26 dpi) or AGI-5198 alone (MS: 24 dpi) treatment groups. These data demonstrate that AGI-5198 does not provide a survival benefit in wtIDH glioma bearing mice (Supplementary Figure 4B).

### Pharmacological inhibition of IDH1-R132H increases PD-L1 expression on mIDH1 Glioma Cells In Vivo

Numerous studies have shown that PD-L1 is highly expressed by wtIDH1 gliomas compared with mIDH1 gliomas (22). We evaluated the PD-L1 expression levels on mIDH1 and wtIDH mouse glioma cells *in vivo*. Mice were either implanted with wtIDH1 or mIDH1 glioma cells and tumors were processed for flow cytometry analysis as indicated in Figure 3A. We observed ∼ 2-fold (*P* ≤ 0.0001) decrease in PD-L1 expression on mIDH1 glioma cells when compared to wtIDH glioma cells (Figure 3B). Next, we evaluated whether AGI-5198 therapy would change the PD-L1 expression on mIDH1 glioma cells *in vivo*. Mice bearing mIDH1 tumors were treated with AGI-5198 as indicated in Figure 3A. Administration of AGI-5198 therapy to mIDH1 glioma bearing mice led to ∼2-fold (*P* ≤ 0.0001) increase in the PD-L1 expression on the tumor cells compared to untreated mIDH1 glioma bearing mice (Figure 3B). The PD-L1 expression on wtIDH1 glioma cells was comparable to mIDH1 glioma treated with AGI-5198 (Figure 3B). Consistent with our results, TCGA analysis of grades II and III glioma patients harboring IDH1-R132H with *TP53* and *ATRX* inactivating mutations revealed ∼1.1-fold (*P* ≤ 0.001) lower expression of PD-L1 (gene annotation denoted as CD274) (23) mRNA compared to wtIDH glioma patients (Figure 3C). We also observed that grades II and III glioma patients harboring IDH1-R132H with 1p/19q have ∼1.1-fold (*P* ≤ 0.001) have lower expression of CD274 mRNA compared wtIDH glioma patients (Figure 3C).

**Figure 3:**
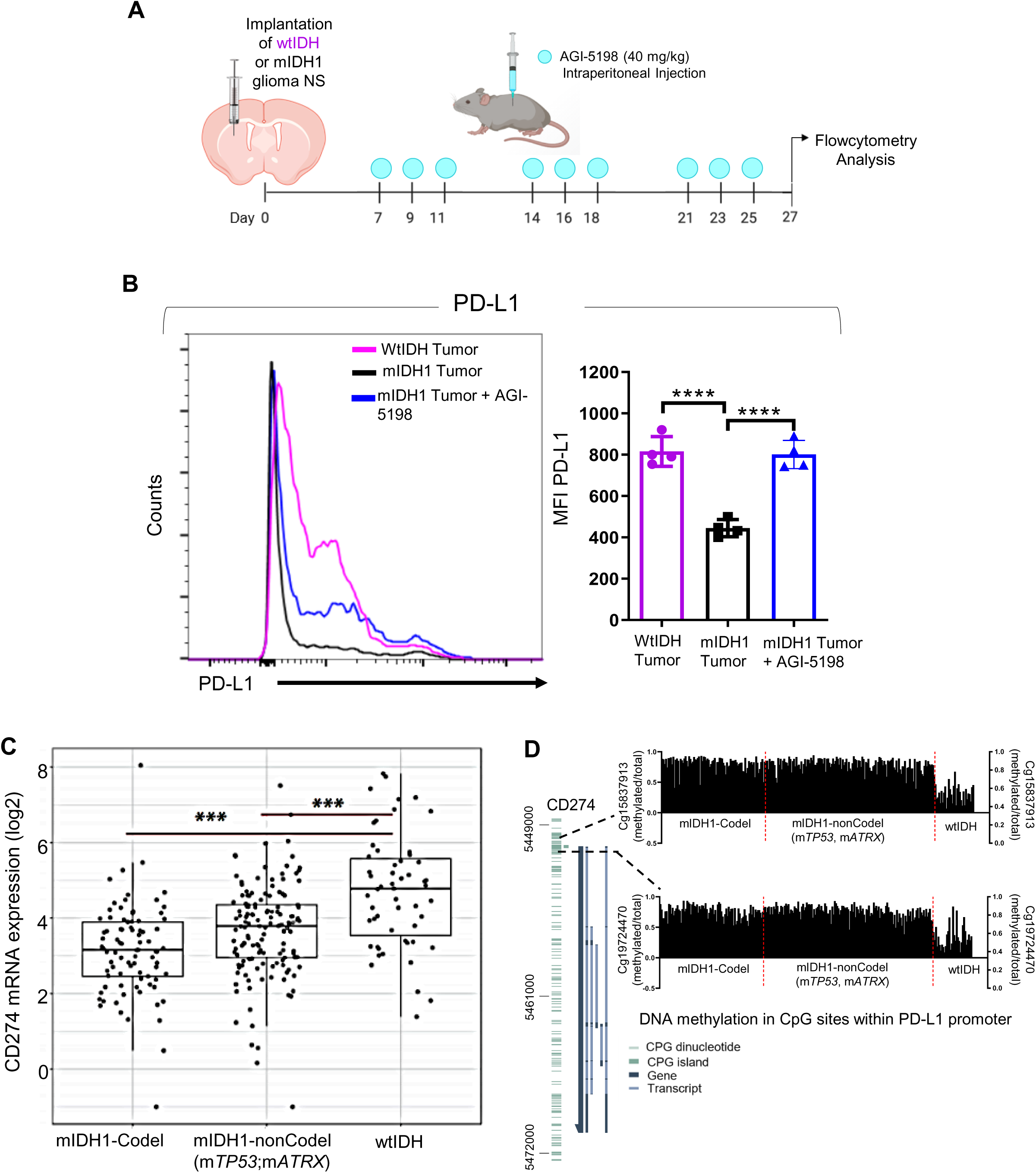
Inhibition of IDH1-R132H increases PD-L1 expression levels on mouse and human mIDH1 gliomas. (A) Diagram of experimental design to characterize PD-L1 expression on glioma cells. At end of the treatment (27 dpi) brains were harvested for flow cytometry analysis. (**B)** Representative histograms display PD-L1 expression levels on CD45^-^/Nestin^+^/Katushka+ glioma cells (purple = wtIDH1 tumor, black = mIDH1 tumor, blue = mIDH1 tumor treated with AGI-5198). \*\*\*\**P* < 0.0001, One-way ANOVA test. Bars represent mean ± SEM (*n* = 4 biological replicates). (**C**) Analysis of CD274 (PD-L1) gene expression for IDHmut-codel (n=85), IDH1mut-noncodel (n=141), and wtIDH (n=55) grade II and III glioma patients. RNA-seq data was obtained from TCGA (GlioVis platform). Graph displays the log_2_ expression value of CD274 mRNA expression: each dot represents 1 patient. \*\*\**P* < 0.001, Tukey’s Honest Significant Difference test. (**D**) DNA methylation levels within cg15837913 and cg19724470 probes in the CpG island of CD274 promoter were determined for IDH mut-codel (n=85), IDH1mut-noncodel (n=141), and wtIDH (n=55) grade II and III glioma patients. Each black bar represents one patient. DNA Methylation status was assessed for TCGA data using Mexpress.

Since 2HG inhibits DNA and histone demethylases (11), altering the epigenome and gene expression patterns, we assessed the CD274 promoter methylation profiles of grades II and III glioma from the TCGA data set using Mexpress. Two probes (cg15837913 and cg19724470) within the CpG island were used to identify DNA methylation within the CD274 promoter. These probes have previously been shown to differ in methylation levels between mutant IDH1 tumor and normal brain tissue (23). We found that DNA methylation of CD274 were higher in both cg15837913 and cg19724470 sites for patients harboring IDH1-R132H with *TP53* and *ATRX* inactivating mutations and patients harboring IDH1-R132H with 1p/19q when compared to wtIDH glioma patients (Figure 3D). These data indicate that IDH-R132H is directly involved in regulating PD-L1 expression in grade II and III gliomas.

### Therapeutic Efficacy of IDH1-R132H inhibition, Standard of Care (SOC) and Anti-PD-L1 blockade in an Intracranial mIDH1 Glioma Model

Given that PD-L1 expression increased on mIDH1 glioma cells when treated with AGI-5198 (Figure 3B), we investigated whether combining AGI-5198 therapy with SOC and anti-PD-L1 (αPD-L1) immune checkpoint blockade would provide a therapeutic benefit to mIDH1 glioma bearing mice. Mice bearing mIDH1 tumors were treated at the indicated doses and treatment schedule specified in Figure 4A. Administration of TMZ alone (MS: 39 dpi), SOC (TMZ+IR) alone (MS: 38 dpi) or SOC+αPD-L1 (MS: 39 dpi) to mice bearing mIDH1 tumor did not confer any survival benefit compared with saline control (MS: 35 dpi; p > 0.05; Supplementary Figure 5). However, administration of αPD-L1 (MS:43 dpi), αPDL+TMZ (MS: 42 dpi), or αPD-L1+ IR (MS: 43 dpi) conferred a ∼1.2-fold (p < 0.01) survival benefit (Supplementary Figure 5).

**Figure 4:**
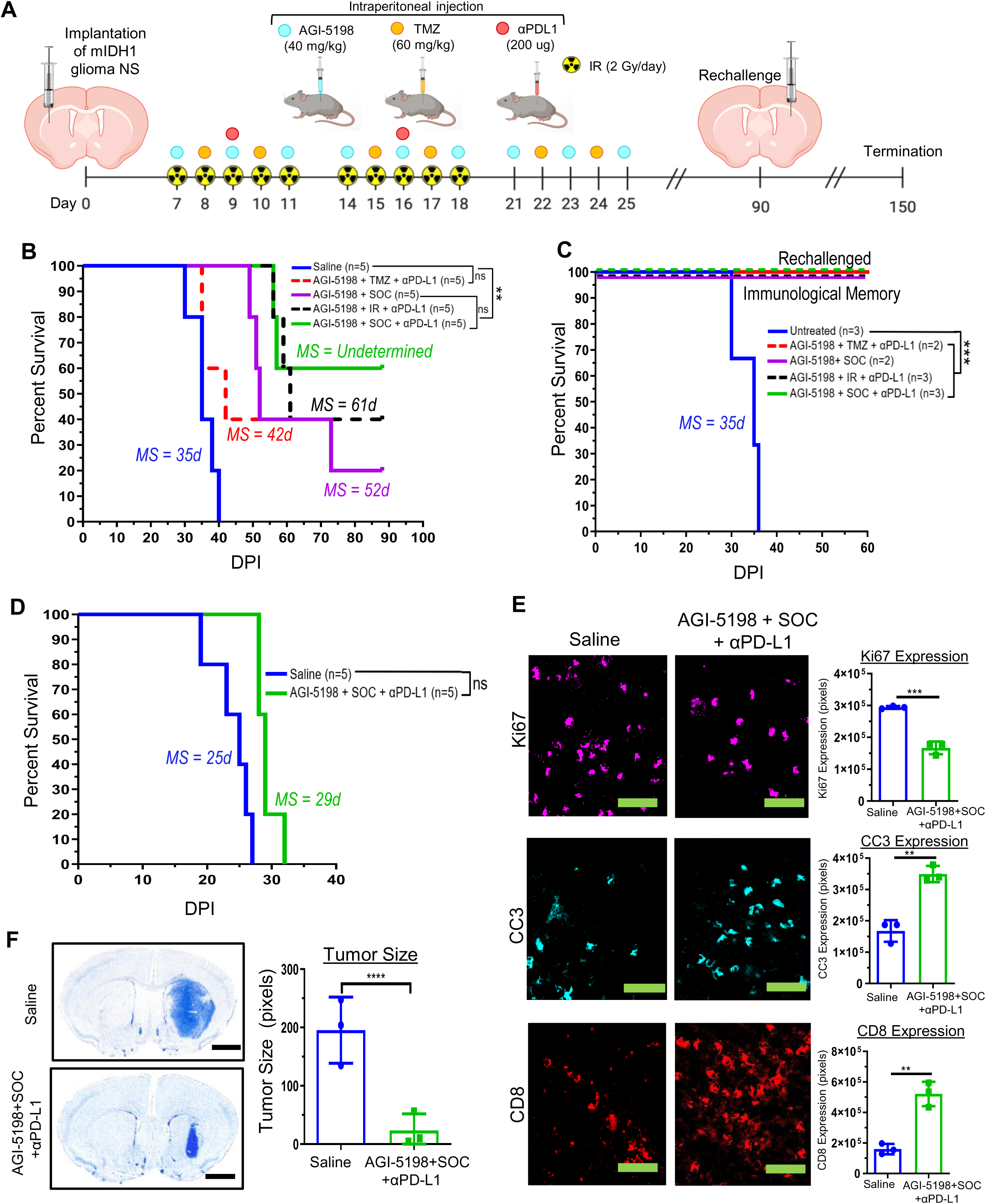
Inhibition of IDH1-R132H in combination with SOC and αPD-L1 immune checkpoint blockade exhibits enhanced survival of mIDH1 glioma bearing mice. (A) Diagram of experimental design to assess survival for AGI-5198 therapy in combination with SOC (TMZ +IR) and αPD-L1 **B)** Kaplan-Meier survival analysis of saline (*n* = 5), AGI-5198 + TMZ + αPD-L1 (*n* = 5), AGI-5198 + SOC (n = 5), AGI-5198 + IR + αPD-L1 (*n* = 5), and AGI-5198 + SOC + αPD-L1 (*n* = 5) treated mice. **(C)** Kaplan-Meier survival plot for rechallenged long-term survivors from (B) AGI-5198 + TMZ + αPD-L1 (*n* = 1), AGI-5198+SOC (*n* = 2), AGI-5198 + IR + αPD-L1 (*n* = 2), and AGI-5198 + SOC + αPD-L1 (*n* = 3), and control mice (*n* = 3). Data were analyzed using the log-rank (Mantel-Cox) test. \*\**P* < 0.001; \*\*\**P* < 0.001; MS = median survival; d = days. (**D**) Kaplan-Meier analysis of saline (*n* = 5) or AGI-5198 + SOC + αPD-L1 (*n* = 5) treated mIDH1 glioma bearing CD8-knockout mice. (**E**) C57BL/6 mice bearing mIDH1 glioma were treated with saline or AGI-5198 + SOC + αPD-L1 as detailed in (A). At 27 dpi, brains were harvested for immunohistochemistry analysis. Immunofluorescence staining for Ki67, cleaved caspase-3 (CC3), and CD8 was performed on 50 µm vibratome tumor sections (green scale bar = 10 μm). Bar graphs represent total number of positive cells for Ki67, CC3, and CD8 in saline or AGI-5198+ SOC+αPD-L1 treatment groups. \*\**P* < 0.01; \*\*\**P* < 0.001, unpaired t-test. Bars represent mean ± SEM (*n* = 3 biological replicates). **(F)** Nissl staining of 50 µm brain sections from saline and AGI-5198+SOC+αPD-L1 treated mIDH1 tumor bearing mice at 27 dpi. \*\*\*\**P* < 0.001; unpaired t-test (black scale bar = 1mm). Bars represent mean ± SEM (*n* = 3 biological replicates).

Treating mIDH1 tumor bearing mice with AGI-5198+SOC (MS: 52) enhanced the efficacy of SOC alone (MS: 38 dpi, Supplementary Figure 4), and resulted in 20% long-term survivors (Figure 4B). We also observed that the addition of AGI-5198 to TMZ+αPD-L1 (MS:42 dpi) or IR+αPD-L1 (MS: 61 dpi) treatments significantly enhanced the efficacy of the therapy for mIDH1 glioma bearing mice (Figure 4B). Notably, there were 40% long-term survivors in AGI-5198+TMZ+αPD-L1 and AGI-5198 +IR+αPD-L1 treatment groups, but the highest survival advantage was observed for AGI-5198+SOC+αPD-L1 (MS: not reached) with 60% long term survivors (Figure 4B; Table 2). The long-term survivors from the AGI-5198+TMZ+αPD-L1, AGI-5198+SOC, AGI-5198+IR+αPD-L1, or AGI-5198+SOC+αPD-L1 treatment groups were rechallenged with mIDH1 tumors in the contralateral hemisphere (Figure 4A) and remained tumor free without further treatment, compared to untreated control mice implanted with tumors which succumbed due to tumor burden (MS: 35 days; *P* ≤ 0.0001) (Figure 4C). These results suggest the development of immunological memory in the tumor bearing animals treated with AGI-5198+TMZ+αPD-L1, AGI-5198+SOC, AGI-5198+IR+αPD-L1, or AGI-5198+ SOC+αPD-L1 treatment groups.

**Table 2:** Log-rank (Mantel-Cox) Test Kaplan Meir Survival Analysis for AGI-5198 treatment in combination with SOC and αPD-L1 immune check point blockade

| Group | P value |
| --- | --- |
| Saline vs. AGI-5198 + TMZ + αPD-L1 | 0.0686 |
| Saline vs. AGI-5198 + SOC | 0.0018 |
| Saline vs. AGI5198 + IR + αPD-L1 | 0.0018 |
| Saline vs. AGI5198 + SOC + αPD-L1 | 0.0018 |
| AGI-5198 + TMZ + αPD-L1 vs. AGI-5198 + SOC | 0.9331 |
| AGI-5198 + TMZ + αPD-L1 vs. AGI5198 + IR + αPD-L1 | 0.6013 |
| AGI-5198 + TMZ + αPD-L1 vs. AGI5198 + SOC + αPD-L1 | 0.3386 |
| AGI-5198 + SOC vs. AGI5198 + IR + αPD-L1 | 0.3655 |
| AGI-5198 + SOC vs. AGI5198 + SOC + αPD-L1 | 0.1599 |
| AGI5198 + IR + αPD-L1 vs. AGI5198 + SOC + αPD-L1 | 0.6943 |

To determine whether the efficacy of AGI-5198+SOC+αPD-L1 is mediated by the adaptive immune system, CD8-knockout mice were implanted with mIDH1 cells and treated with saline or AGI-5198+SOC+αPD-L1 at the indicated dose and treatment schedule (Figure 4A). There was not a statistically significant difference in MS between saline or AGI-5198+SOC+αPD-L1 treated mice. These data indicate the critical role that CD8^+^ T cells have in mediating the observed therapeutic response induced by AGI-5198+SOC+αPD-L1 treatment (Figure 4D).

Subsequently, we analyzed the *in vivo* expression of Ki67 (proliferation marker), Cleaved Caspase 3 (CC3; apoptosis marker), and CD8 (cytotoxic T cell marker) in mIDH1 tumors, two days after saline, or AGI-5198+SOC+αPD-L1 treatments (27 dpi, Figure 4A). Combining AGI-5198 treatment with SOC and αPD-L1 resulted in a significant decrease in Ki67 expression (*P* < 0.0001) and an increase in CC3 expression (*P* < 0.01) compared to the saline treated group (Figure 4E). We also observed increased infiltration of CD8^+^ T cells (*P* < 0.01) in tumors treated with AGI-5198+SOC+αPD-L1 when compared to saline. Additionally, we quantified the size of the tumor two days after AGI-5198+SOC+αPD-L1 or saline treatment (27 dpi). We observed a ∼3-fold (*P* < 0.0001) decrease in tumor size in the AGI-5198+SOC+αPD-L1 treated mice (Figure 4F).

We also assessed complete blood cell counts (CBC), serum biochemistry for aminotransferase (ALT), bilirubin, urea (BUN) and creatinine for mice treated with saline or AGI-5198+SOC+αPD-L1 two days following the last treatment dose (Figure 4A). No differences were observed in relation to CBC between the treatment groups (Supplementary Figure 6). The levels of ALT, BUN and creatinine were within the normal range, indicating normal functioning liver and renal systems (Table 3). Furthermore, liver tissue sections from both treatment groups showed no signs of necrosis, inflammation, or changes in cellular structures (Supplementary Figure 7).

**Table 3:** Mice serum levels of biochemical variables after intraperitoneal treatment with AGI-5198 + IR+ TMZ + αPDL1 (n=3).

| Group | Creatinine (μmol/L) | BUN (mmol/L) | ALT (U/L) | AST (U/L) |
| --- | --- | --- | --- | --- |
| Saline | 0.11 ± 0.17 | 30 ± 1.7 | 73 ± 7.2 | 303 ± 290.6 |
| AGI5198 + IR+TMZ+<br>αPDL1 | 0.21 ± 0.02 | 19.3 ± 6.7 | 112.3 ± 31 | 311 ± 213.6 |

**Table 4.** Flowcytometry Antibodies

| Antibodies | Source | Catalog Number |
| --- | --- | --- |
| V450 rat anti mouse CD45 (clone: 30-F11), Flow 1:200 | BD Bioscience | Cat# 560501 |
| PE Rat anti mouse F4/80 (clone: BM8), Flow 1:200 | Biolegend | Cat# 123110 |
| Alexa Fluor 700 Rat anti mouse CD206 (clone: C068C2), Flow 1:200 | Biolegend | Cat# 141733 |
| PE armenian hamster anti mouse CD11c (clone: N418), Flow 1:200 | Biolegend | Cat#117308 |
| PercpCy5.5 rat anti mouse B220 (clone: RA36B2), Flow 1:200 | Biolegend | Cat #: 103236 |
| FITC rat anti mouse CD8 (clone: KT15) , Flow 1:1:00 | Thermofisher | Cat # MA5-16759 |
| PercpCy5.5 armenian hamster anti mouse CD3 (clone: 145-2C11), Flow 1:200 | Biolegend | Cat # 100328 |
| FITC rat anti mouse Gr-1 (clone: RB6-8C5), Flow 1:200 | Biolegend | Cat # 108405 |
| PercpCy5.5 rat anti mouse CD11b (clone: M1/70), Flow 1:200 | Biolegend | Cat # 101230 |
| FITC rat anti mouse CD4 (clone:GK1.5) , Flow 1:200 | Biolegend | Cat # 100405 |
| APC rat anti mouse CD25 (clone: 3C7), Flow 1:200 | Biolegend | Cat # 101909 |
| APC rat anti mouse CD44 (clone: IM7), Flow 1:200 | Biolegend | Cat #: 103012 |
| APC/Cy rat anti mouse CD62L (clone: MEL-14), Flow 1:200 | Biolegend | Cat #: 104428 |
| PE rat anti mouse IFNγ (clone: XMG1.2), Flow 1:200 | Biolegend | Cat #: 505808 |
| PE rat anti mouse PD-L1 (clone: B7-H1), Flow 1:200 | Biolegend | Cat #: 124308 |
| APC/Cy rat anti mouse PD-1 (clone: 29F.1A12), Flow 1:200 | Biolegend | Cat #: 135224 |
| APC rat anti mouse TIM 3 (clone: B8.2C12), Flow 1:200 | Biolegend | Cat #: 134008 |
| PE anti mouse TIGT(clone: 1G9), Flow 1:200 | Biolegend | Cat #: 142104 |
| PE/Cy7 armanian hamster anti mouse CD80 (clone: 16-10A1), Flow 1:200 | Biolegend | Cat #: 104734 |
| APC/Cy anti mouse CD86 (clone: GL-1), Flow 1:200 | Biolegend | Cat #: 105030 |
| FITC anti mouse MHC II (clone: 39-10-8) , Flow 1:200 | Biolegend | Cat #: 115006 |
| FITC rat anti mouse Foxp3 (clone MF-14), Flow 1:200 | Biolegend | Cat # 126403 |
| PE-Tetramer, Flow 1:100 | MBL<br>International | Cat # TB-5001-1 |
| Alexa Fluor 647 rabbit anti mouse Calreticulin (clone: EPR3924) , Flow 1:50 | Abcam | Cat # ab196159 |
| Alexa Fluor 647 rabbit ant mouse pH3Ser10 (clone: D2C8), Flow 1:50 | Cell Signaling | Cat # 3458S |
| Efluor 780-Live/Dead, Flow 1:200 | Affymetrics | Cat #: 65-0865-14 |
| Annexin V/PI Apoptosis Kit | Thermofisher | Cat #: V13241 |
| Click-iT EdU Kit | Thermofisher | Cat #: C10418 |
| Mouse Foxp3 Buffer Set | BD Biosciences | Cat #: 560409 |
| BD CytoFix/Cytoperm Kit | BD Biosciences | Cat #: 554714 |

**Table 5.** Immunohistochemistry Primary Antibodies

| **Antibodies** | **Source** | **Catalog Number** |
| --- | --- | --- |
| Rat anti mouse MBP, IHC 1:300 | Millipore | Cat# MAB386 |
| Rabbit anti mouse CD8, IHC: 1: 2000 | Cedarlane | Cat #: 361003 |
| Rat anti mouse GFAP, IHC: 1:1000 | Millipore | Cat #: AB5541 |
| Rabbit anti mouse Cleaved Caspase 3, IHC: 1:1000 | Sell Signaling | Cat # 9661S |
| Rabbit anti mouse Ki-67 | Abcam | Cat # ab15580 |

**Table 6.**
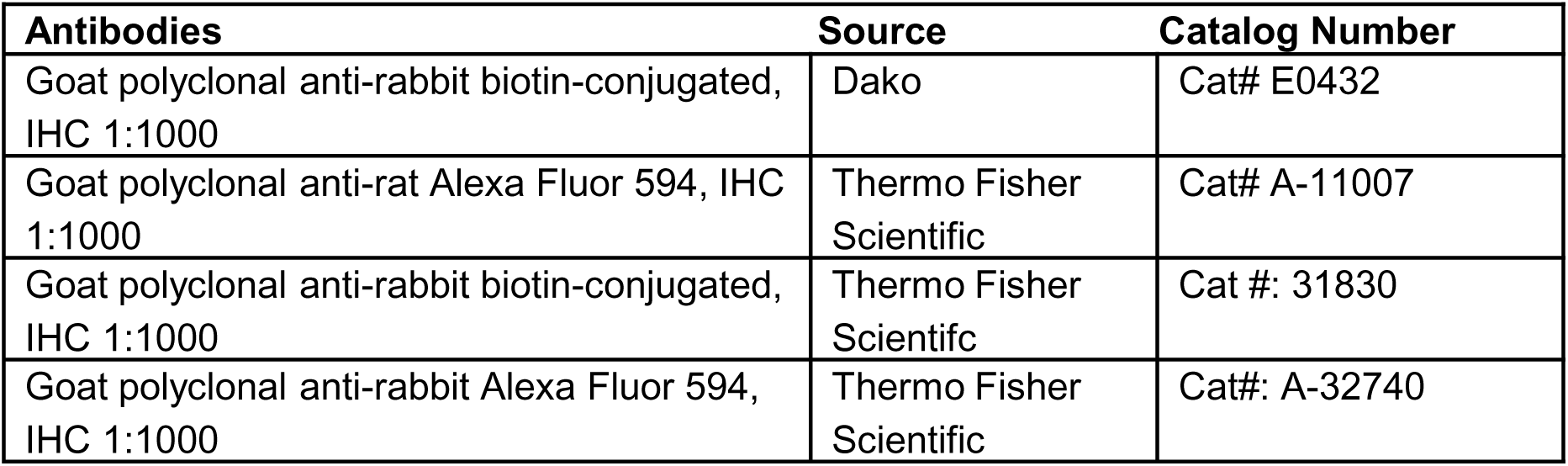
Immunohistochemistry Secondary Antibodies

| **Antibodies** | **Source** | **Catalog Number** |
| --- | --- | --- |
| Goat polyclonal anti-rabbit biotin-conjugated, IHC 1:1000 | Dako | Cat# E0432 |
| Goat polyclonal anti-rat Alexa Fluor 594, IHC 1:1000 | Thermo Fisher Scientific | Cat# A-11007 |
| Goat polyclonal anti-rabbit biotin-conjugated, IHC 1:1000 | Thermo Fisher Scientific | Cat #: 31830 |
| Goat polyclonal anti-rabbit Alexa Fluor 594, IHC 1:1000 | Thermo Fisher Scientific | Cat#: A-32740 |

**Table 7.** Reagents for In vivo Studies

| **Reagent** | **Source** | **Catalog Number** |
| --- | --- | --- |
| Polyethylene Glycol 400 | Sigma-Aldrich | Cat#: 8170035000 |
| AGI-5198 | Amatek Chemicals | Cat#:1355326-35-0 |
| Temozolomide | Selleckchem | Cat #: S1237 |
| Rat Anti-mouse PD-L1 | BioXcell | Cat# BE0101 |
| Ethanol | Sigma-Aldrich | Cat #: 459836 |

Long-term survivors from the AGI-5198+SOC+αPD-L1 treatment group rechallenged with mIDH1 glioma were euthanized after 60 days (Figure 4A). Potential disruptions of brain tissue caused by the therapy were assessed by immunohistochemical staining for myelin basic protein (MBP) as a marker for myelin sheaths integrity and glial fibrillary acid protein (GFAP) as marker of astrocyte integrity. We did not observe demyelination or changes in astrocyte integrity in mIDH1 glioma rechallenged mice from AGI-5198+SOC+αPD-L1 treatment group compared to the saline treated mice (Supplemental Figure 8). These data demonstrate that the AGI-5198+SOC+αPDL treatment does not induce changes in brain architecture in the long-term survivors. Overall, our data demonstrate that administration of AGI-5198+SOC+αPD-L1 treatments confers an overall survival advantage and elicits an anti-mIDH1 glioma immune response with no overt acute or long-term off-target toxicity.

### Expansion of mIDH1 Anti-Glioma Specific Cytotoxic T Cells in Response to IDH1-R132H inhibition in Combination with SOC and αPD-L1 Immune Checkpoint Blockade

Given that AGI-5198 therapy in combination with SOC and αPD-L1 immune checkpoint blockade resulted in tumor regression and long-term survival in mIDH1 glioma bearing mice (Figure 4B, C), we aimed to study the anti-mIDH1 glioma specific immune response elicited by this treatment strategy. Mice bearing mIDH1 tumors harboring a surrogate tumor antigen, ovalbumin (OVA) were treated with saline, AGI-5198+SOC, or AGI-5198+SOC+αPD-L1 as indicated in Figure 5A. Mice were euthanized two days following the last treatment dose and brains were processed for flow cytometry analysis to characterize the immune cell infiltration in the TME. Treatment of mIDH1 glioma bearing mice with AGI-5198+SOC therapy induced a ∼1.6-fold (*P* ≤ 0.05) increase in the percentage of tumor infiltrating plasmacytoid DCs (pDCs: CD45^+^/CD11c^+^/B220^+^) and ∼2.2-fold (*P* ≤ 0.001) increase conventional DCs (cDCs: CD45^+^/CD11c^+^/B220^-^) compared to saline treated mice (Figure 5B). In mice treated with AGI-5198+SOC+αPD-L1, we observed an additional ∼1.5-fold (*P* ≤ 0.05; Figure 5B) increase in tumor infiltrating pDCs and ∼1.3-fold (P ≤ 0.001; Figure 5B) increase in tumor infiltrating cDCs compared to mice treated with AGI-5198+SOC.

**Figure 5:**
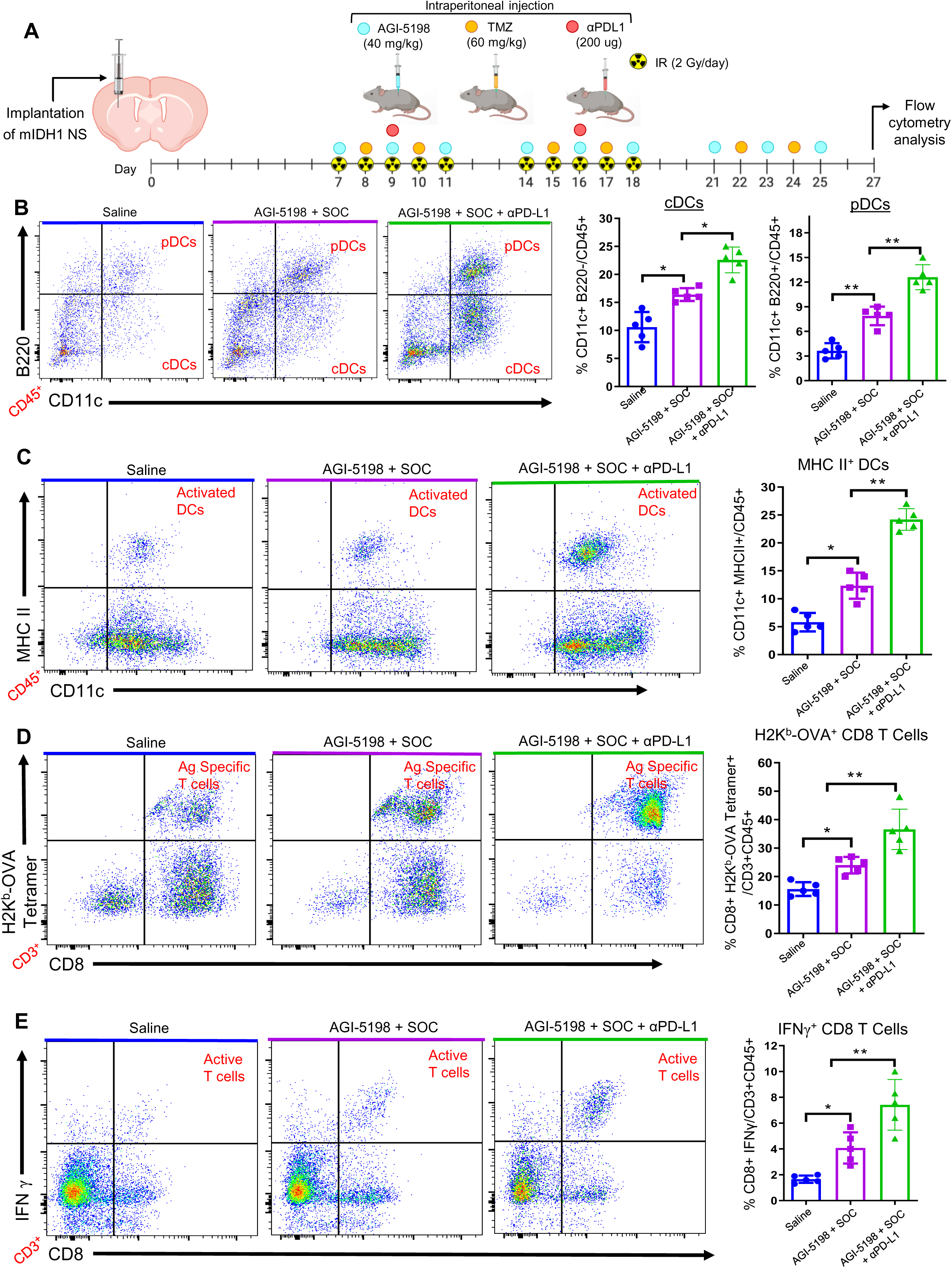
Inhibition of IDH1-R132H in combination with SOC and αPD-L1 immune checkpoint blockade induced tumor-specific CD8 T cell responses within mIDH1 glioma TME. (A) Diagram of experimental design to characterize immune cellular infiltrates in the TME of mIDH1 glioma bearing mice after treatment with saline, AGI-5198 + SOC or AGI-5198 + SOC + αPD-L1 therapies. **(B)** The percent of pDCs (CD11c^+^/B220^+^) and cDCs (CD11c^+^/B200^-^) within the CD45^+^ cell population in the TME of saline, AGI-5198+IR, or AGI-5198 + SOC + αPD-L1 treated mIDH1 tumor bearing mice was assessed at day 27 dpi. Representative flow plots for each group are displayed. \**P* < 0.05; \*\**P* < 0.01, one-way ANOVA test. Bars represent mean ± SEM (*n* = 5 biological replicates). **(C)** Activation status of CD11c^+^ DCs in the mIDH1 TME was assessed by expression of MHC II. Representative flow plots for each treatment are displayed. \**P* < 0.05; \*\*\**P* < 0.001, one-way ANNOVA test. Bars represent mean ± SEM (*n* = 5 biological replicates). **(D)** Tumor-specific CD8^+^ T cells within the TME of mIDH1-OVA tumors were analyzed by staining with the SIINFEKL-K^b^ tetramer. Representative flow plots for each treatment are displayed. \*\**P* < 0.01; \*\*\**P* < 0.001, one-way ANNOVA test. (**E**) Activation status of CD8^+^ T cells within the TME of mIDH1-OVA tumors was analyzed by staining for IFNγ after TME stimulation with the tumor cell lysate. \*\**P* < 0.01; \*\*\**P* < 0.001, one-way ANOVA test. Bars represent mean ± SEM (*n*= 5 biological replicates).

To determine the effect AGI-5198+SOC+αPD-L1 treatment on DC activation status, we measured expression levels of co-stimulatory molecules CD80, CD86 and MHC II (24–26) in the DCs of the mIDH1 TME. We observed an increase in MHC II (∼2.4 fold, *P* ≤ 0.05), CD80 (0.8 fold, *P* ≥ 0.05), and CD86 (∼2.0 fold, *P* ≤ 0.05) CD45^+^/CD11c^+^ DCs positive cells in the TME of AGI-5198+SOC treated mice when compared to saline treated mice (Figure 5C, Supplementary Figure 9). The activation status of DCs was further enhanced by the addition of αPD-L1 blockade to AGI-5198+SOC treatment. We observed an increase in the frequency of MHC II (∼2.0-fold, *P* ≤ 0.01), CD80 (∼2.1-fold, *P* ≤ 0.01), and CD86 (∼1.8-fold, *P* ≤ 0.001) DCs in the TME (Figure 5C, Supplementary Figure 8). These data suggest that AGI-5198+SOC+αPD-L1 treatment promotes an enhanced adaptive immune response within the mIDH1 glioma TME through activation of antigen presenting DCs.

To further evaluate the specificity of the anti-mIDH1 glioma immune response elicited by AGI-5198+SOC+αPD-L1 treatment, we assessed antigen specific CD8^+^ T cells responses in the TME of mIDH1-OVA glioma bearing mice in each treatment group. The mIDH1-OVA model allowed us to monitor the generation of glioma specific T cells and to detect them by using the SIINFEKL-H2K^b^ tetramer. We observed a ∼1.6 fold (*P* ≤ 0.05) increase in tumor antigen-specific CD3^+^/ CD8^+^/ SIINFEKL-H2K^b^ tetramer^+^ T cells in the TME of mIDH1-OVA tumor bearing mice treated with AGI-5198+SOC compared to saline treatment group (Figure 5D). This was also enhanced in mice treated with AGI-5198+SOC+αPD-L1 blockade by ∼1.4 fold (*P* ≤ 0.01; Figure 5D). Blood serum analysis showed that interleukin-12 (IL12), a cytokine critical for CD8^+^ cytotoxic cell activity (27), increased ∼1.6-fold (*P* ≤ 0.001) and ∼2.7-fold (*P* ≤ 0.001) in mice treated with AGI-5198+SOC+αPD-L1 compared to AGI-5198+SOC or saline treated mice, respectively (Supplemental figure 10).

We also examined the impact of AGI-5198+SOC and AGI-5198+SOC+αPD-L1 therapy on the activation status of CD3^+^/CD8^+^ T cells in the TME through quantification of interferon-γ (IFNγ) expression levels by flow cytometry. The IFNγ levels of CD8^+^ T cells isolated from the TME of mice treated with AGI-5198+SOC were ∼2 fold (*P* ≤ 0.05) greater than those of CD8^+^ T cells isolated from control mice treated with saline (Figure 5E). This response also increased ∼1.4 fold (*P* ≤ 0.01) in mice treated with AGI-5198+SOC+αPD-L1 (Figure 5E). Collectively, these data demonstrate that AGI-5198+SOC+αPD-L1 therapy mediates a robust anti-tumor immune response through the expansion and activation of mIDH1 glioma-specific CD8^+^ T cells.

### Treatment of mIDH1-glioma Bearing Mice with IDH1-R132H inhibition in combination with SOC and Anti-PD-L1 Blockade Decreases the Presence of Immunosuppressive Cells in the TME

Gliomas have been shown to employ numerous mechanisms to suppress the immune system (28); e.g., accumulation of immunosuppressive cells, such as myeloid-derived suppressor cells (MDSCs), regulatory T cells (Tregs), and tumor-associated M2 macrophages (MØs) which inhibit CD8^+^ T cell activation and expansion (28). We evaluated if AGI-5198, SOC, and αPD-L1 blockade treatment reduces the accumulation of immunosuppressive MDSCs, Tregs, and M2 macrophages (MØ) in the mIDH1 glioma TME. Mice with mIDH1-OVA tumors were treated with saline, AGI-5198+SOC, or AGI-5198+SOC+αPD-L1. Brains were processed for flow cytometry analysis at 27 dpi as indicated in Figure 6A. We observed a 1.4-fold (*P* ≤ 0.05) decrease in the percentage of tumor infiltrating MDSCs (CD45^+^/CD11b^+^/Gr-1^+^) in the AGI-5198+SOC treated mice compared to the saline (Figure 6B). We also observed ∼2.3 fold (*P* ≤ 0.001; Figure 6B) decrease in tumor infiltrating MDSCs in animals treated with AGI-5198+SOC+αPD-L1 compared to mice treated with AGI-5198+SOC. Flow cytometry analysis for Tregs (CD45^+^/CD4^+^/CD25^+^/Foxp3^+^) also showed a decrease in mIDH1 glioma by∼ 2.4-fold (*P* ≤ 0.05; Figure 6C) in mice treated with AGI-5198+SOC compared to saline, and an additional decrease of ∼1.5 fold (*P* ≤ 0.05) in mice treated with AGI-519+SOC+αPD-L1 (Figure 6C). Moreover, we observed a ∼1.7 fold (*P* ≤ 0.0001) decrease in the number of tumor infiltrating M2 MØs (CD45^+^/F480^+^/CD206^+^) the TME of AGI-5198+SOC compared to saline treated mice, and a ∼1.5-fold (*P* ≤ 0.05) decrease in mice treated with AGI-519+SOC+αPD-L1 (Figure 6D). Conversely, we observed a ∼1.5 fold (*P* ≤ 0.001) increase in the number of M1 MØs in the TME of AGI-5198+SOC+αPD-L1 treated mice compared to mIDH1 glioma bearing mice treated with AGI-5198+SOC or saline (Figure 6D). Overall, we observed that AGI-5198+SOC+αPD-L1 treatment significantly decreases the amount of immunosuppressive MDSCs, Tregs, and M2 MØs in the TME of mIDH1 glioma bearing mice compared to AGI-5198+SOC and saline treatment groups.

**Figure 6:**
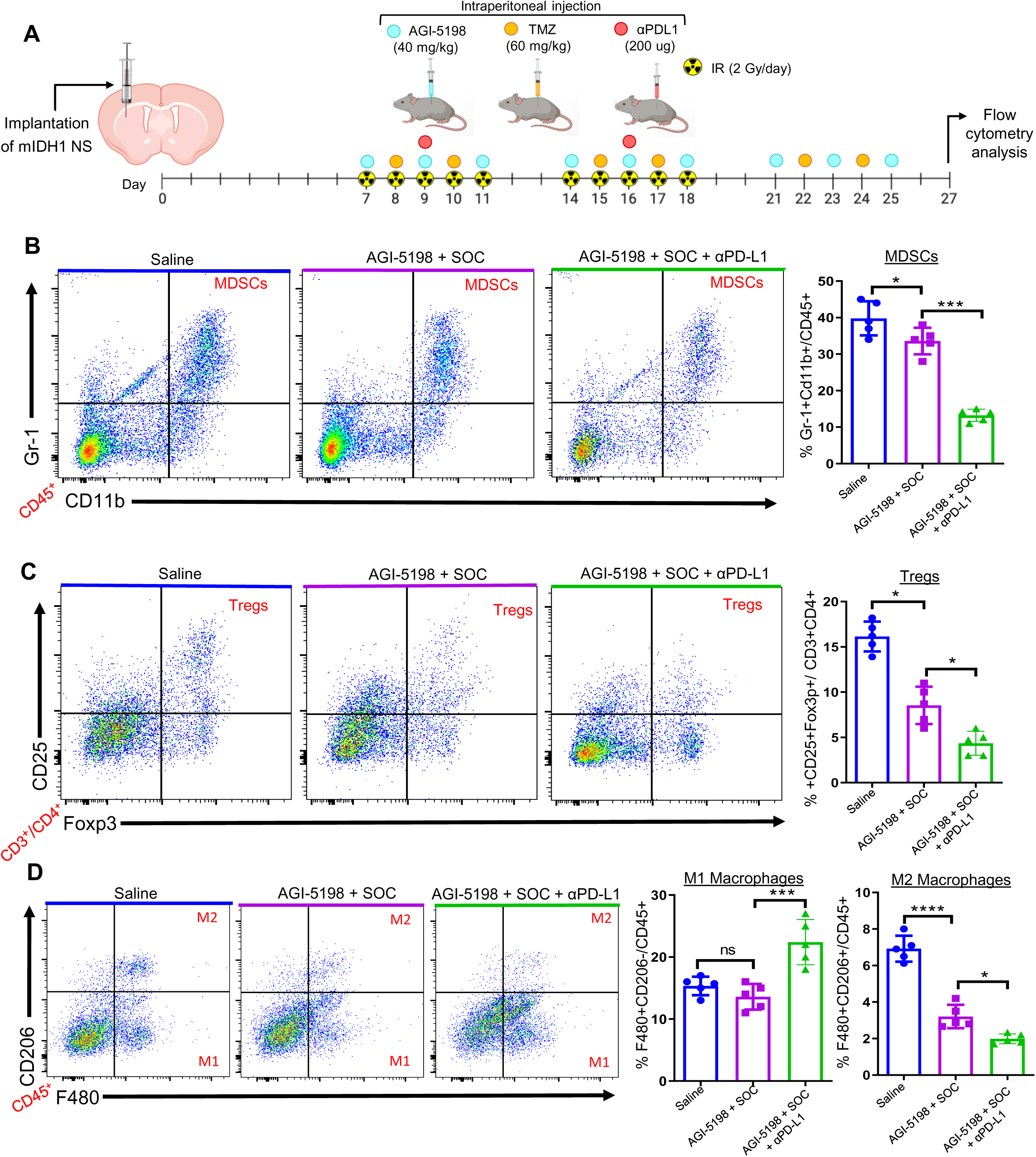
Inhibition of IDH1-R132H in combination with SOC and αPD-L1 immune checkpoint blockade decreased the accumulation of immunosuppressive cells in the TME of mIDH1 glioma. (A) Diagram of experimental design to characterize immune cell infiltration in the TME of mIDH1 glioma bearing mice after treatment with saline, AGI-5198 + SOC or AGI-5198 + SOC + αPD-L1 therapies. **(B)** The percent of MDSCs (CD11b^+^/Gr1^+^) within the CD45^+^ cell population in the TME of saline, AGI-5198 + IR, or AGI-5198 + SOC + αPD-L1 treated mIDH1 glioma bearing mice was assessed at 27 dpi. Representative flow plots for each group are displayed. \**P* < 0.05; \*\*\**P* < 0.001, one-way ANOVA test. Bars represent mean ± SEM (*n* = 5 biological replicates). **(C)** The percent of regulatory T cells (Tregs, CD4^+^/CD25^+^/Foxp3^+^) within the CD45^+^ cell population in the TME of saline, AGI-5198 + IR, or AGI-5198 + SOC + αPD-L1 treated mIDH1 glioma bearing mice was assessed at 27 dpi. Representative flow plots for each group are displayed. \**P* < 0.05, one-way ANOVA test. Bars represent mean ± SEM (*n* = 5 biological replicates). **(D)** The percent of M1(F480^+^/CD206^+^) and M2 (F480^+^/CD206^+^) macrophages within the CD45^+^ cell population in the TME of saline, AGI-5198 + IR, or AGI-5198 + SOC + αPD-L1 treated glioma bearing mice was assessed at 27 dpi. Representative flow plots for each group are displayed. \**P* < 0.05; \*\*\*\**P* < 0.0001, one-way ANOVA test. Bars represent mean M1 and M2 macrophage quantification ± SEM (*n* = 5 biological replicates).

### Impact of oncometabolite 2HG on T Cell Functions

D-2HG is produced by mIDH1 glioma neurospheres (7) and released into the mIDH1 glioma TME. Thus, we next wished to evaluate the impact of D-2HG on T cell functions. Numerous studies have shown D-2HG to be poorly cell-permeable (22). We examined the effect of D-2HG on the activation and proliferation status of antigen specific CD8 T cells from OT-1 transgenic mice, which have T cell receptors engineered to recognize the SIINFEKL peptide (19). OT-1 splenocytes were fluorescently labeled with 5-(and-6)-carboxyfluorescein diacetate succinimidyl ester (CFSE) and stimulated for four days with 100nM SIINFEKL peptide in the presence or absence of 0.25µM D-2HG (Figure 7A). Proliferation and activation of T cells was assessed by CFSE dilution analysis and quantification of IFNγ expression levels in the supernatants, respectively. Unstimulated OT-1 T cells did not proliferate and showed low IFNγ levels (Figure 7B-D). However, almost 100% of T cells from OT-1 mice underwent antigen-specific proliferation in response to SIINFEKL stimulation and IFNγ levels in the supernatant were ∼5-fold higher than unstimulated splenocytes (Figure 7B-D). The addition of D-2HG to stimulated T cells did not significantly alter proliferation of IFNγ levels compared to cells treated with SIINFEKL alone (Figure 7B-D). These data indicate that D-2HG does not directly impact T cell activation or proliferation.

**Figure 7:**
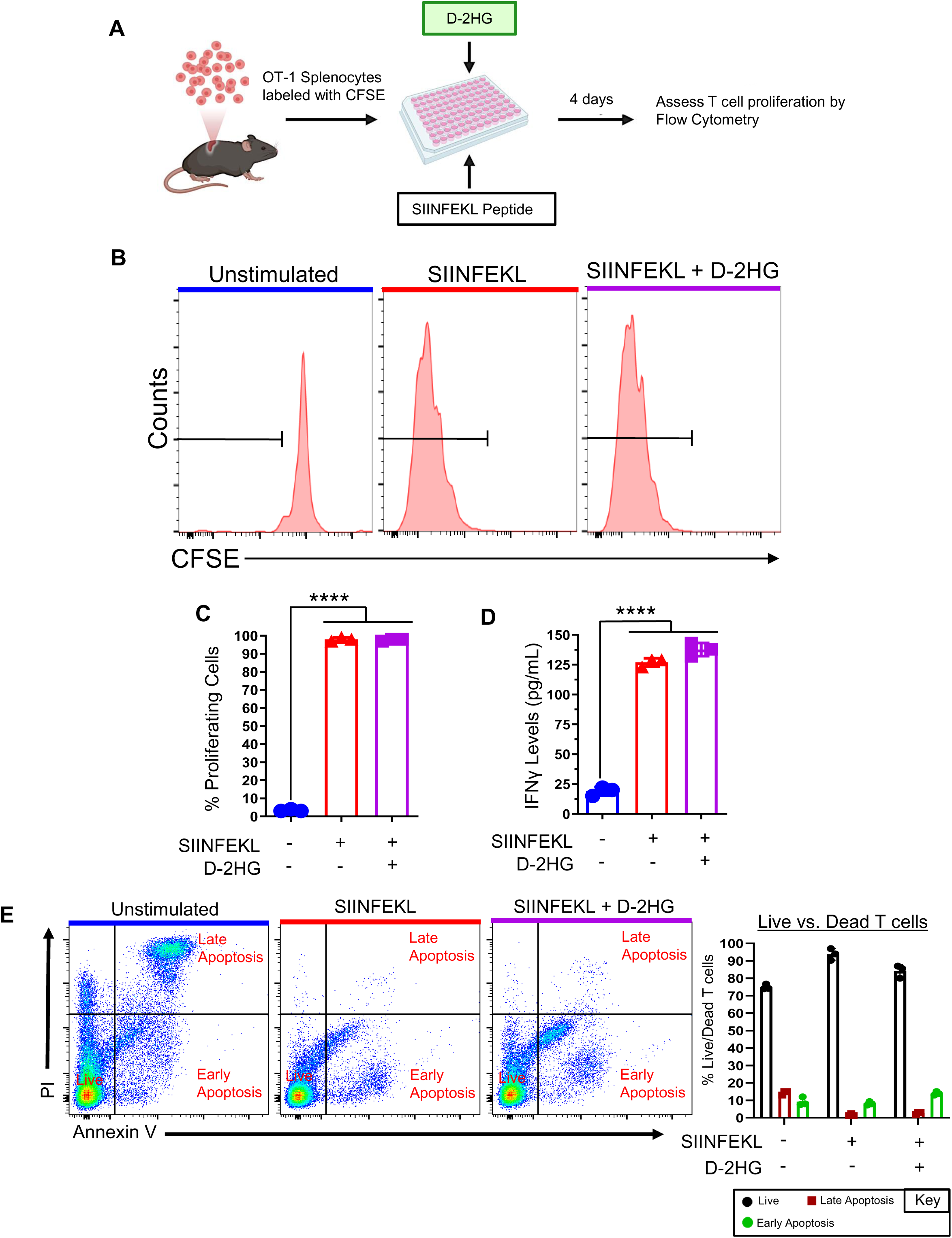
D-2HG does not inhibit T cell proliferation and activation. (A) Diagram of experimental design to analyze 2HG dependent inhibition of T cell proliferation. (**B**) Flow plots show representative CFSE stains of unstimulated splenocytes (inactivated T cells), splenocytes undergoing proliferation in response to 100nM SIINFEKL (activated T cells), and the effect of 25µM D-2HG on SIINFEKL-induced T cell proliferation. (**C**) Quantification of OT-1 splenocytes undergoing T cell proliferation. \*\*\**P* < 0.001, one-way ANOVA test. Bars represent mean ± SEM (*n* = 3 technical replicates). (**D**) Quantification of IFNγ levels in the supernatants of OT-1 splenocytes stimulated with SIINFEKL in the presence of D-2HG. IFNγ levels were assessed by ELISA. **(E)** Representative flow plots of OT-1 splenocytes incubated with 25µM D-2HG in the presence of 100 nM SIINFEKL peptide for 4 days and stained with Annexin V-FITC and Propidium iodide (PI). Live activated T cells (CD3^+^/CD8^+^/IFNγ^+^) were identified as Annexin V negative and PI negative. Dead cells undergoing early apoptosis were identified as Annexin V positive and PI negative. Dead T cells undergoing late apoptosis were identified as Annexin V positive and PI positive. Quantitative analysis of the distribution of live and dead cells during 4-day incubation period with D-2HG.

Since T cell functions require viable T cells, we next evaluated whether D-2HG would induce apoptosis of activated T cells. OT-1 splenocytes were stimulated with 100nM SIINFEKL peptide in the presence or absence of 0.25µM D-2HG for four days (Figure 7A). Activated T cells (CD3^+^/CD8^+^/IFNγ^+^) undergoing early or late apoptosis were identified by flow cytometry with AnnexinV/PI staining. In accordance with our data on T cell activation and proliferation, we observed ∼90% live (Annexin V^-^/ PI^-^) T cells in culture with SIINFEKL alone compared to ∼76% live T cells in cultures with unstimulated OT-1 cells (Figure 7E). The addition of D-2HG to the T cell culture with SIINFEKL did not induce apoptosis (Figure 7E). Notably, when non-activated T cells were cultured with 15mM and 50mM D-2HG, we observed apoptosis in greater than ∼40% cells (Supplementary Figure 11). These data indicate that at higher doses D-2HG is toxic for CD8 T cells, rather than inhibiting their proliferating capacity. Overall, our data indicate that D-2HG, which is present in the mIDH1 glioma TME, does not alter T cell functions.

### IDH1-R132H Inhibition in Combination with SOC and Anti-PD-L1 Immune Checkpoint Blockade Confers Memory T Cell Response Following Glioma Rechallenge

We demonstrated that mIDH1 glioma bearing mice survive long-term when treated with AGI-5198+SOC+αPD-L1 immune checkpoint blockade and that long-term survivors exhibit immunological resistance to mIDH1 glioma rechallenge in the contralateral hemisphere (Figure 3B, C). Therefore, we assessed the T cell memory response in the rechallenged long-term survivors treated with AGI-5198+SOC+αPD-L1 (Figure 8A). Normal untreated mice implanted with mIDH1 glioma were utilized as controls. Mice were euthanized fourteen days post-rechallenge, brains and blood were processed for flow cytometry analysis of CD45^+^/CD3^+^/CD8^+^/CD44^[high]^/CD62L^[low]^ effector (T_EM_) and CD45^+^/CD3^+^/CD8^+^/CD44^[high]^/ CD62L^[high]^ central (T_CM_) memory T cells. Mice treated with AGI-5198+SOC+αPD-L1 showed an increase in the percentage of TME infiltrating T_EM_ (∼8-fold; *P* ≤ 0.0001) and T_CM_ (∼3.3-fold; *P* ≤ 0.0001) cells compared to the untreated control mice (Figure 8B). We also observed an increase in the percentage of T_EM_ (∼1.6-fold; *P* ≤ 0.0001) and T_CM_ (∼7-fold; *P* ≤ 0.0001) cells in the blood of mice treated with AGI-5198+SOC+αPD-L1 compared to the control (Figure 8C). Blood serum analysis showed that interleukin-15 (IL15), a cytokine critical for CD8^+^ memory T cell activity (29), increased ∼4-fold (*P* ≤ 0.001) in mice treated with AGI-5198+SOC+αPD-L1 compared to untreated mice (Supplemental figure 12).

**Figure 8:**
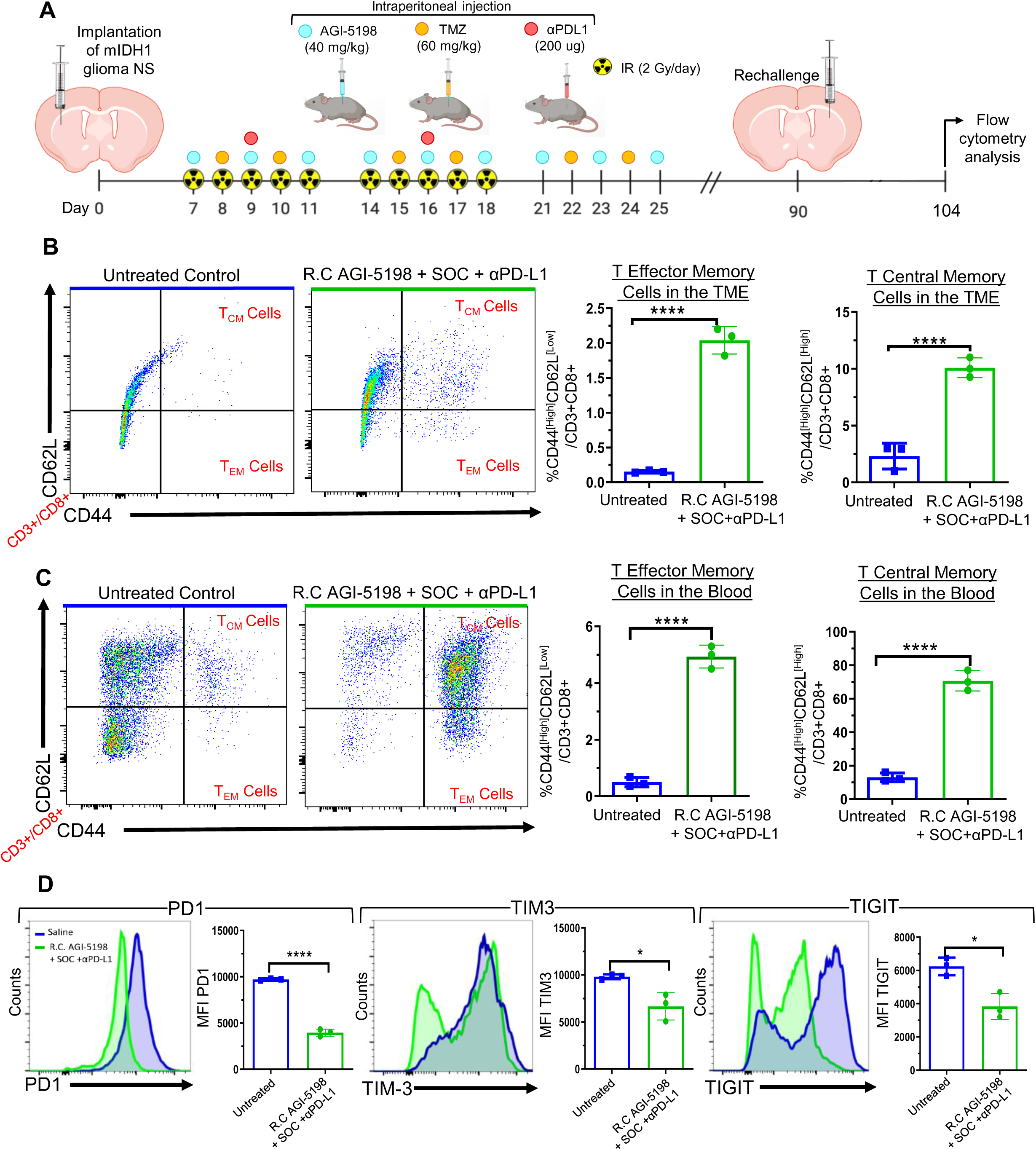
Inhibition of IDH1-R132H in combination with SOC and αPD-L1 immune checkpoint blockade induces CD8 T cell memory responses and lowers T cell exhaustion within the TME of mIDH1 glioma bearing mice. (A) Diagram of experimental design to assess memory T cell response in mIDH1 glioma bearing mice treated with AGI-5918 in combination with SOC and anti-PD-L1 immune checkpoint blockade. **(B)** The percent of T central memory cells (T_CM_ cells) (CD62L^+^/CD44^+^) and T effector memory cells (T_EM_ cells) (CD62L^-^/CD44) within the CD45^+^/CD3^+^ cell population in the TME of untreated control and AGI-5198 + SOC + αPD-L1 treated mice was assessed at 7 dpi after mIDH1 glioma rechallenge. Representative flow plots for each group are displayed. \*\*\*\**P* < 0.0001, unpaired t-test. Bars represent mean ± SEM (*n* = 3 biological replicates). **(C)** The percent of T central memory cells (CD62L^+^/CD44^+^) and T effector memory cells (CD62L^-^/CD44) within the CD45^+^/CD3^+^ cell population in the circulation of untreated control and AGI-5198 + SOC + αPD-L1 treated mice was assessed at 7 dpi after mIDH1 glioma rechallenge. Representative flow plots for each group are displayed. \*\*\*\**P* < 0.0001, unpaired t-test. Bars represent mean ± SEM (*n* = 3 biological replicates). **(D)** CD8+ T cell exhaustion in the TME of mIDH1 glioma bearing mice of untreated control and AGI-5198 + SOC + αPD-L1 treated mice was assessed at 14 days post tumor rechallenge, by assessing the expression levels of PD-1, TIM-3, and TIGIT markers. Representative histograms display each marker’s expression levels (blue = saline, green =AGI-5198 + SOC + αPD-L1). \**P* < 0.05; \*\*\*\**P* < 0.0001, unpaired t-test. Bars represent mean ± SEM (*n* = 3 biological replicates).

Since αPD-L1 immune checkpoint blockade may augment the effector functions of glioma infiltrating cytotoxic T cells, we tested the effect of AGI-5198+SOC+αPD-L1 treatment on T cell exhaustion by measuring the expression levels of PD-1, TIM-3, and TIGIT markers (30) on the CD8 T cells in the TME of rechallenged mIDH1 glioma bearing mice. We observed a decrease in PD-1 (∼3.3 fold, *P* ≤ 0.001), TIM-3 (∼1.3-fold, *P* ≤ 0.05), and TIGIT (∼0.6 fold, *P* ≤ 0.05) expression on CD45^+^/CD3^+^/CD8^+^ T cells in the TME of rechallenged AGI-5198+SOC+αPD-L1 treated mice compared to untreated control mice (Figure 8D). These data indicate that the combination of AGI-5198, SOC and αPD-L1 treatment reduces T-cell exhaustion and strongly favors the generation of a memory CD8^+^ T cell response against mIDH1 glioma leading to the generation of adaptive immune response against tumor recurrence.

## Discussion

A critical factor influencing the success of therapeutic approaches for gliomas expressing IDH1-R132H in the context of *TP53* and *ATRX* inactivation (mIDH1) is the ability to harness memory T cells’-mediated response in order to prevent tumor recurrence. Recently there has been substantial focus on utilizing small molecule inhibitors to target the gain of function enzymatic activity of IDH1-R132H (17). Several preclinical studies have shown that small molecule inhibitors targeting IDH1-R132H have been effective in impairing tumor progression when used as monotherapy in pre-clinical models of acute myeloid leukemia (AML) and glioma (15, 17, 22, 31, 32).

We have previously shown that the genetic context in which IDH-R132H mutation occurs impacts tumor biology and treatment outcomes (7). For insistence, we demonstrated that D-2HG epigenetically reprograms the expression of genes involved in DNA-damage response in gliomas harboring IDH1-R132H in the context of ATRX and TP53 inactivation (mIDH1), eliciting glioma radioresistance (7). Treatment of mouse and human mIDH1 glioma cells with IDH1-R132H inhibitor *in vitro*, radiosensitized mIDH1 glioma cells. Another study has also demonstrated that IDH1 gliomas radioresistance was determined by the genetic context in which IDH-R132H was expressed (33). These studies suggest that the response to therapies for gliomas harboring mutant IDH1 is dependent on the genetic context in which this mutation is found (7, 33, 34). Also, ATRX inactivation can have profound effects on chromatin structure, which in turn will affect the transcriptional profile of the cells in which this genetic lesion is encountered (35–37).

A critical feature of any proposed therapeutic strategy for mIDH1 gliomas should be the ability to prevent tumor recurrence, which is a salient clinical feature of this disease (38, 39). Immunological memory after immunotherapy, mediated by tumor-specific memory T cells can protect against tumor recurrence (40). A previous study demonstrated that in primary IDH1-R132H grade II gliomas there is less infiltration of CD8^+^ T cells in the TME; CD8^+^ T cells were predominantly localized in the perivascular niche (41). This reduction in T cells’ numbers was due to decreased expression of cell adhesion molecules (ICAM1) and chemo-attractants (CXCL9 and CXCL10), which mediate the recruitment of T cells from circulation into the TME (41). Similarly, another study demonstrated less intratumoral infiltration of CD8+ T cells in syngeneic GL261 and Sleeping Beauty derived SB28 (driven by NRAS, PDGFβ, and short hairpin-targeting TP53) models expressing IDH1-R132H compared to wildtype IDH1 tumors. The authors demonstrated reduced IFN-γ expression in the T cells present within the IDH1-R132H glioma TME (22). Further, reduced expression of genes involved in leukocyte migration was observed in IDH1-R132H tumor-bearing Ntva_Ink4a/Arf +/- mice (6). However, none of these studies addressed IDH1-R132H in the genetic context of ATRX and TP53 inactivation.

In this study, we aimed to assess the immunological impact of blocking 2-HG production on anti-mIDH1 glioma immunity in a genetically engineered mutant IDH1 mouse glioma model, which also encodes for ATRX and TP53 loss, recapitulating the salient genetic lesions encountered in mutant IDH1 astrocytomas (2, 7). We demonstrate that systemic administration of an IDH1-R132H inhibitor prolonged the MS of mIDH1 glioma bearing mice from 34 days to 43 days (*P* < 0.05), with 40% long-term survivors (Figure 2). Furthermore, when the long-term survivors from the IDH1-R132H inhibition treatment group were rechallenged with mIDH1 tumor cells in the contralateral hemisphere, all the mice remained tumor free without further treatment, indicating the development of anti-glioma immunological memory.

A recent study suggests that the D-2HG directly inhibits anti-glioma CD8 T cells’ functions in mIDH1 glioma TME (42). However, previous reports have shown that D-2HG is poorly cell-permeable (9, 43). Herein, we demonstrate that D-2HG does not suppress the functions of mouse antigen-specific CD8 T cells’ proliferation (Figure 7). Moreover, D-2HG induced apoptosis in >40% of T cells at 15 mM and 50 mM (Supplementary Figure 10), indicating that at the doses previously reported D-2HG might have been toxic for CD8 T cells, rather than inhibiting their proliferation (44, 45).

Immunogenic cell death (ICD) elicited by dying tumor cells contributes to the development of adaptive immunity (46). Our results show that blocking 2HG production in mouse and human mIDH1 glioma increases the expression of CRT, the release of ATP and HMGB-1 DAMPs molecules (Figure 1). Although, we have shown that mIDH1 mouse and human glioma cells are radioresistant (7) (Figure 2), we observed that mIDH1 cells treated with IR alone released similar levels of ATP as IDH1-R32H inhibition treatment group (Figure 1). In this respect, it has been previously shown that tumor cells treated with radiotherapy release high levels of ATP, triggering autophagy activation (47–49). Thus, to tolerate IR induced stress, the mIDH1 mouse and human glioma cells could be releasing elevated levels of ATP to activate the autophagy pathway for survival.

ICD elicits the activation of antigen presenting DCs to prime cytotoxic CD8^+^ T cells and leads to a robust anti-mIDH1 glioma response in our model. Mice bearing mIDH/ATRX KD/TP53 KD glioma treated with IDH1-R132H inhibitor in combination with SOC demonstrated an increase in pDCs and cDCs in the TME compared to the saline treated group. This treatment strategy also upregulated the expression of co-stimulatory ligands CD80, CD86 and MHC II on DCs (24–26), polarizing them towards an activated phenotype (Figure 5, Supplementary Figure 8). In addition, using a mIDH1 mouse glioma model harboring surrogate tumor antigen, ovalbumin (OVA), we monitored the generation of mIDH1 glioma specific T cells in the TME using the H2K^b^-tetramer. Our data show an increase in mIDH1-glioma specific T cells in the TME of mIDH1 glioma bearing mice treated with IDH1-R132H inhibitor and SOC compared to saline treated mice (Figure 5). We also observed a significant decrease in the accumulation of immunosuppressive MDSCs, Tregs, and M2 macrophages in the TME of mIDH1 glioma bearing mice after inhibition of 2HG production combined with SOC vs. saline treated mice (Figure 6). These data strongly suggest that inhibiting IDH-R132H in combination with SOC induces a mIDH1 glioma-specific immune response.

Immunotherapy has emerged as a powerful tool to address cancer eradication, but it fails to elicit a potent or lasting response in the clinical arena (28). This has been partly attributed to the presence of inhibitory receptor molecules on T cells, such as programmed cell death protein 1 (PD-1) (28). A study evaluating the immunological gene profile of 282 primary IDH1-R132H gliomas compared to 151 wildtype IDH gliomas from TCGA, determined that the gene and protein expression of PD-L1 was significantly lower in mutant IDH1 gliomas compared to wildtype IDH1 glioma (23). We analyzed the expression of PD-L1 in TCGA data set of wildtype IDH, IDH1-R132H with 1p/19q codeletion, and IDH1-R132H with ATRX/TP53 mutation in grades II and III gliomas. The expression of IDH-R132H was associated with lower levels of PD-L1 expression in glioma patients with 1p/19q codeletion and ATRX/TP53 mutations (Figure 3). TCGA analysis also revealed that IDH1-R132H increased DNA methylation at two CpG sites within the PD-L1 promoter in 1p/19q co-deletion and 1p/19q non-codeletion grade II and III gliomas. This assertion is also supported by *Mu et al* (23). IDH-R132H induced epigenetic reprogramming of the PD-L1 gene locus could increase DNA methylation at the CpG sites within the PD-L1 promoter. In this study, we demonstrate that genetically engineered mIDH1 mouse glioma, which recapitulates the salient genetic lesions encountered in mutant IDH1 astrocytomas exhibits lower levels of PD-L1 compared to wtIDH1 mouse glioma (Figure 3). We observed that inhibiting IDH-R132H increases the PD-L1 expression on mIDH1 glioma cells in the TME by ∼ 2-fold compared to untreated control mice (Figure 3). Since higher expression of PD-L1 on tumor cells leads to T cell exhaustion (30) and negatively regulate T cell activation, priming and expansion (50) we decided to combine inhibition of 2HG production, with immune check point blockade using a αPD-L1 blocking antibody. The efficacy of PD-1/PD-L1 immune checkpoint blockade has not been investigated in in glioma models harboring IDH1-R132H in the context of ATRX and TP53 loss. We observed that PD-L1 immune checkpoint blockade when used as monotherapy, elicited a small increase in MS of mice bearing mIDH1 glioma, with no long-term survivors (Supplementary Figure 4). There is evidence that shows that immune-checkpoint blockade used as monotherapy has failed in Phase III clinical trials to improve overall survival of patients with glioma (20). Our data demonstrate that co-administering αPD-L1 immune checkpoint blockade with inhibition of 2HG production and SOC significantly improved the MS of mIDH1 glioma bearing mice (Figure 4). We also observed a reduction of immunosuppressive cells, i.e, MDSCs, Tregs, and M2 macrophages in the TME of mIDH1 glioma bearing mice treated with IDH1-R132H inhibition in combination with SOC and αPD-L1 compared to saline treated control mice (Figure 6). Also, this strategy reduced T cell exhaustion and favored the generation of memory CD8 + T cells, which elicited a strong immunological memory response against mIDH1 glioma rechallenge (Figure 8).

The role of memory T cells in promoting immunological memory after therapy is critical for preventing tumor recurrence. Our proposed strategy consists of targeting the mIDH1 glioma TME with a combination of D-2HG inhibition, SOC, and αPD-L1 immune checkpoint blockade. We show that this treatment is capable of prolonging animal survival as a result of an increase in tumor-specific infiltrating CD8 T cells and memory T cells, which are needed to prevent tumor recurrence (Figure 4-6,8). In summary, our pre-clinical data provides evidence that support the further development of combining IDH1-R132H inhibition, with SOC and αPD-L1 in a Phase I clinical trial for glioma patients expressing IDH1-R132H in context of TP53, ATRX inactivating mutations.

## Materials and Methods

Additional details on the methods used in this study are provided in the Supplementary Materials.

### Study Approval

All animal experimental studies were performed in compliance with Institutional Animal Care & Use Committee (IACUC) at the University of Michigan.

### Mouse glioma models

Genetically engineered mouse models that recapitulate the salient features of human glioma are critical to understanding the mechanisms of tumor progression and response to therapeutics (7, 35). Our laboratory used the Sleeping Beauty Transposon System to generate genetically engineered immunocompetent wtIDH and mIDH1 mouse models of glioma harboring ATRX and TP53 loss. The wtIDH glioma includes genetic lesions in NRASG12V, *Atrx* and *Tp53* knockdown (35). The mIDH1 glioma model includes genetic lesions in NRASG12V, *Atrx* and *Tp53* knockdown with the addition of IDH1-R132H (7). Mouse neurospheres (NS) derived from the genetically engineered mouse glioma models were utilized to perform experiments in this study.

### Statistical analysis

Sample sizes were selected based on preliminary data from pilot experiments and previously published results in the literature and our laboratory. All animal studies were performed after randomization. Unpaired Student t-test or one-way analysis of variance (ANOVA), followed by Tukey’s multiple comparisons post-test were utilized for comparing experimental groups with controls from immunofluorescence, tumor size quantification, flow cytometry analysis and T cell functional assays. Turkey’s Honest Significant Difference test was utilized to comparing glioma patient groups from TCGA. Log-rank (Mantel-Cox) test analysis was utilized to compare treatment groups from Kaplan-Meier survival curves. Data were analyzed using Prism 6.0 (GraphPad Software). Data were normally distributed so that the variance between groups was similar. P values less than 0.05 were considered statistically significant. All values are reported as means ± SD with the indicated sample size. No samples were excluded from analysis.

## Author Contributions

P.K., S.V.C., J.C.G., M.B.G-F., F.J.N., F.M.N., Y.L., M.Y., D.L., and M.B.E., performed experiments; P.K., J.C.G, F.J.N., M.S.A., Y.L, A.S., P.R.L., and M.G.C., analyzed the data; P.K., J.C.G., Y.L., A.S, P.R.L., and M.G.C., designed the figures; P.K., S.V.C., J.C.G, M.G.F., F.J.N, F.M.N., Y.L., J.J.M., A.S., P.R.L., and M.G.C., designed the research and contributed to writing and editing the manuscript.

## Acknowledgments

This work was supported by the National Institutes of Health/National Institute of Neurological Disorders & Stroke (NIH/NINDS) Grants R21-NS091555 to M.G.C., A.S. and PRL; R37-NS094804, and R01-NS074387 to M.G.C.; R01-NS076991, R01-NS082311, and R01-NS096756 to P.R.L; National Institutes of Health/National Cancer Institute (NIH/NIC) Grant T32-0CA009676 to M.G.C; National institutes of Health/National Institute of Biomedical Imaging and Bioengineering (NIH/NIBIB) Grant R01-EB022563 to J.J.M., P.R.L, and M.G.C.; University of Michigan M-Cube;the Department of Neurosurgery; the University of Michigan Rogel Comprehensive Cancer Center; Leah’s Happy Hearts Foundation; and the Biointerfaces Institute at the University of Michigan. Experimental design schematics were made using www.biorender.com.

## Supplementary Methods

### Reagents

DMEM-F12, DMEM RPMI-1650, FBS, PBS, N2, and B27 supplements and penicillin-streptomycin were purchased from GIBCO, Life Technologies. Epidermal growth factor (EGF) and fibroblast growth factor (FGF) were purchased from Peprotech. Anti-mouse CD45, CD11c, B220, F4/80, CD206, CD3, CD4, CD8, CD25, CD80, CD86, MHC II, CD44, CD62L, Gr1, CD11b, and PDL1 antibodies for flow cytometry analysis were obtained from Biolegend. SIINFEKL tetramers were obtained from MBL International (Supplementary Table 2). For immunohistochemistry, anti-mouse MBP and GFAP primary antibody was purchased from Millipore; anti-mouse Cleaved Caspase 3 was purchased from Cell Signaling; anti-mouse Ki-67 and Calreticulin were purchased from Abcam; and anti-mouse CD8 was purchased from Cedarlane (Supplementary Table 3). Secondary antibodies for immunohistochemistry were purchased from Thermofisher and Dako (Supplementary Table 4). AGI-5198 compound was purchased from Amatek Chemical Co; Temozolomide was purchased from Selleckchem; and Anti mouse PD-L1 for in vivo studies was purchased from BioXcell (Supplementary Table 5).

### Cell Culture

Mouse wtIDH1 and mIDH1 mouse neurospheres were grown in DMEM-F12 media supplemented with 100 units/mL Penicillin-Streptomycin, 1x B-27, 1x N-2, 100ug/mL Normocin, 20ng/mL hFGF and 20ng/mL hEGF (NSC media). Mouse mIDH1-OVA neurospheres were grown in NSC media supplemented with 100 units/mL penicillin. Human SJGBM2 neurosphere cells were grown in NSC media. Human MGG119 neurosphere cell were grown in Neurobasal media supplemented with 100 units/mL Penicillin-Streptomycin, 1x B-27, 1x N-2, 100ug/mL Normocin, 20ng/mL hFGF and 20ng/mL hEGF. Human SJGBM2 glioma cells were shared by Children’s Oncology Group (COG) Repository, Health Science Center, Texas Tech University; these cells were grown in IMDM media supplemented with 20% fetal bovine serum and 100 units/mL penicillin. MGG119 cells were shared by Dr. Daniel P. Cahill laboratory, Harvard Medical School (1). Neurospheres were dissociated with Accutase detachment reagent when they had to passaged. Cells were maintained in a humidified incubator at 95% air/5% CO2 at 37°C and passaged every 2-4 days.

### Animal Strains

Six to eight-week-old female C57BL/6 and CD8 knockout mice were purchased from Jackson Laboratory (Bar Harbor, ME) and were housed in pathogen free conditions at the University of Michigan.

### Clonogenic Assay

Mouse and human (SJGBM2:wtIDH1, MGG119: mIDH1) glioma cells were seeded at density of 1.0 x 10^6^ cells into 25-cm^2^ flasks containing DMEM medium supplemented with 10% fetal bovine serum, and 100U/mL Penicillin-Streptomycin. Mouse cells were treated with 1.5 μM AGI-5198; human glioma cells were treated with 5µM AGI-5198. DMSO was utilized as vehicle control. Cells were maintained in culture for ten days, media containing AGI-5198 or vehicle control was replaced every 2 days. On day ten, mouse and human glioma cells were plated (10 × 10^4^ cells/well) into 6 well plates in DMEM medium supplemented with 10% fetal bovine serum and 100U/mL Penicillin-Streptomycin. After 24 hours, cells were treated with escalating doses of radiation (Mouse cells: 0, 2, 4, and 8 Gy; Human cells: 0, 5, 10, and 20Gy). Cells were then trypsinized, serially diluted and seeded (cell numbers ranging from 100 to 2000 cells per cell type were plated depending on the radiation dose) in triplicates into 6-well plates. Cells were maintained in a humidified incubator at 95% air/5% CO2 at 37°C for 10 days. Once the colony formation was observed, plates were washed with PBS and cell colonies were stained with 0.25% crystal violet. Colonies were counted using a bright field microscope. The survival fraction was calculated relative to DMSO vehicle control.

### DAMPs Measurement

Mouse mIDH1 neurospheres and MGG119 glioma cells were seeded at density of 1.0 x 10^6^ cells into 25-cm^2^ flasks containing NSC media. Mouse cells were treated with 1.5 μM AGI-5198; human glioma cells were treated with 5µM AGI-5198. DMSO was utilized as vehicle control. Cells were maintained in culture for ten days, media containing AGI-5198 or vehicle control was replaced every 2 days. On day ten, mouse and human glioma cells were plated (10 × 10^4^ cells/well) into 6 well plates in NSC media. After 24 hours, mouse neurospheres and human glioma cells were treated with 3Gy and 10Gy radiation respectively. After an additional 24 hrs, mouse neurospheres and human glioma cells were treated with 1.5µM and 5µM AGI-5198 or DMSO vehicle control respectively. Release of DAMPs was assessed 72 hours post radiation and 48 hours post AGI-5198 treatment. Neurospheres were collected, dissociated with Acqutase and stained with Calreticulin (1:50) in PBS containing 2% FCS (flow buffer) for 25 minutes at 4°C. Cells were then washed three times with flow buffer and the fluorescence of the samples was read on a BD FACS ARIA SORP (BD Bioscience) using the 647 laser (APC setting). Data were analyzed with Flowjo v.10 Software. Levels of HMGB-1, IL-1α, and IL-6 in the cultures’ supernatants was measured by ELISA according to manufacturer’s instructions (R&D) at the Cancer Center Immunology Core, University of Michigan. Levels of ATP in the cultures’ supernatants was measured by ENLITEN® ATP Assay according to manufacturer’s instructions (Promega).

### Determination of 2-HG in brain samples

2HG concentration in the tumor microenvironment of untreated normal mice, wtIDH1 and mIDH1 glioma bearing mice; and mIDH1 glioma bearing mice treated with AGI-5198 was assessed by liquid chromatography-mass spectrometry (UPLC-MS). AGI-5198 (40 mg/kg) was injected intraperitonially (i.p) into mice bearing mIDH1 glioma at 7, 9, 11, 14, 16, 18, 21, 23, 25 days post tumor cell implantation (dpi). Brains from untreated control mice, wtIDH1 and mIDH1 glioma bearing +/- AGI-5198 treatment were harvested for analysis at 27 dpi. Brain tissue (10-15mg) was mixed with 10 μL of internal standard solution containing 10 μg/mL of 2-HG-D3 (Sigma-Aldrich), and was homogenized in 1 mL of methanol/water (80:20, v/v). The homogenate was centrifuged at 12000 rpm for 10 min at 4 °C, and the supernatant was collected and dried under a stream of nitrogen at 37 °C. Then 160 μL of N-(p-toluenesulfonyl)-L-phenylalanyl chloride (TSPC, 2.5 mM in acetonitrile) and 2 μL of pyridine was added, and the samples were incubated at 37 °C for 20 min to derivatize D-2HG. After derivatization the mixture was dried with nitrogen at 37 °C and reconstituted in 100 μL of acetonitrile/water (50:50, v/v). The samples were then centrifuged at 12000 RPM for 10 min and the supernatant was collected for quantification.

The quantification of derivatized 2-HG was carried out using ultra performance UPLC-MS (Waters ACQUITY system). The mobile phase consisted of deionized water containing 0.1% formic acid (A) and acetonitrile/methanol (1:1, v/v) containing 0.1% formic acid (B). The gradient started from 70% A and maintained for 1 min, changed to 30% A over 3 min and maintained for 2 min, and finally changed back to 70% A over 0.5 min and held for 1.5 min. The flow rate was 0.5 mL/min, and column temperature was set at 40 °C. The analysis was performed on a Waters ACQUITY UPLC HSS T3 column (1.8 μm, 3.0 × 75 mm) with mass detection at 448.17 (-) for derivatized 2-HG, and at 451.19 (-) for derivatized 2-HG-D3.

### Intracranial mIDH1 glioma models

Syngeneic tumors were established in C57BL/6 mice by stereotactically injecting 50,000 mIDH1, wtIDH1 or mIDH1-OVA neurospheres into the right striatum using a 22-gauge Hamilton syringe (1 μL over 1 minute) with the following coordinates: +1.00 mm anterior, 2.5 mm lateral, and 3.00 mm deep (2).

### Preparation of AGI-5198 Formulation for *in vivo* Studies

Approximately 5 mg AGI-5198 was dissolved in 350 µl ethanol and 350 µL PEG-400, then 300µL water was added to achieve the final volume of 1 ml (final AGI-5198 concentration in the solution was 5 mg/ml). For a 40 mg/kg intraperitoneal dose, ∼200µL formulation was injected per mouse.

### Radiotherapy

Seven days’ post wtIDH1 or mIDH1 tumor cells’ implantation, a dose of 2 Gy Irradiation (IR) was administered to mice 5 days a week for two weeks. Mice bearing tumor were placed under a copper Orthovoltage source, their body was shielded with iron collimators to focus the irradiation energy beam to the brain (2, 3). Irradiation treatment was given to mice at the University of Michigan Radiation Oncology Core.

### In vivo Cell Cycle Analysis

AGI-5198 (40 mg/kg) or saline was injected i.p into mice bearing mIDH1 glioma at 7, 9, 11, 14, 16, 18, 21, 23, 25 days post tumor cell implantation (dpi). A course of IR (2 Gy) was administered to mice 5 days a week for two weeks starting at 7 dpi (2, 3). At 27 dpi, a dose of 10 mg/kg EdU was i.p injected into mIDH1 tumor bearing mice 3 hours before they were sacrificed. Tumors were dissected and made into single cell suspensions. Immune cells were labeled with CD45 magnetic beads (Miltenyi) following the manufacturer’s instructions. After 15 min of incubation at 4 °C, cells were washed and passed through a preconditioned MS column placed in the magnetic field of a suitable MACS Separator. Tumor cells (CD45 negative) were collected and resuspended in flow buffer. Tumor cells were fixed and permeabilized using the BD kit (BD Biosciences) and stained with pH3-Ser10 (1: 50; Cell Signaling). Click-reaction was performed to detect EdU positive cells following the manufacturer’s instructions (ThermoFisher) (2, 8).

### Therapeutic Studies in Tumor Bearing Animals

AGI-5198 (40 mg/kg) or saline was injected i.p into mice bearing mIDH1 or wtIDH1 glioma at 7, 9, 11, 14, 16, 18, 21, 23, 25 dpi. A course of IR (2 Gy) was administered to mice 5 days a week for two weeks starting at 7 dpi (2, 3). To assess the therapeutic efficacy of AGI-5198 in combination with standard of care (IR + Temozolomide) and anti-PDL1 immune checkpoint blockade mice harboring mIDH1 glioma were injected i.p with AGI-5198 (40 mg/kg) or saline on days 7, 11, 14, 16 18, 21, 23, and 25 dpi. Temozolomide (TMZ: 60 mg/kg) was administered to mice i.p. on days 8, 10, 15, 17, 22, and 24 dpi. A course of IR (2 Gy) was administered to mice 5 days a week for two weeks starting at 7 dpi. Mice were injected i.p with αPD-L1 (8 mg/kg) at 9 and 16 dpi. For survival studies, mice that displayed signs of tumor burden were perfused with Tyrode’s solution and paraformaldehyde (PFA). For phenotypic characterization of immune cellular infiltrates, mice were processed for flow cytometry analysis at 27 dpi. For immunological memory response assessment, long-term survivors were rechallenged with mIDH1 glioma cells in the contralateral hemisphere at 90 dpi and processed for flow cytometry analysis at 104 dpi; or the experiment was terminated 60 dpi post tumor rechallenge.

### Immunohistochemistry

For immunofluorescence assessment, brain fixed in PFA were serially sectioned 50 µm thick using the vibratome system (Leica VT100S). Sections were placed consecutively into six wells (12-well plate) containing 2 mL of PBS with 0.01% sodium azide. Each well containing sections was a representation the whole brain. For neuropathology assessment, brains fixed in PFA were embedded in paraffin and sectioned 5 µm thick using a microtome system (Leica RM2165. Immunohistochemistry was performed on brain sections by permeabilizing them with TBS-0.5% Triton-X (TBS-TX) for 20 min. This was followed by antigen retrieval at 96 °C with 10 mM sodium citrate (pH 6) for an additional 20 min. Once the sections cooled down to room temperature (RT), they were washed five times (3min per wash) with TBS-TX and blocked with 10% horse serum in TBS-TX for 1 hour at RT. Brain sections were incubated in primary antibody Ki-67 (1:1000), cleaved caspase 3 (1:400), GFAP (1:1000), MBP (1:500), or CD8 (1:2000) diluted in 1% horse serum TBS-TX overnight at RT. The next day sections were washed with TBS-TX 5 times. Secondary antibodies were diluted 1:1000 in 1% goat serum TBS-TX. Brain sections labeled with Ki67, CC3, CD8 were incubated in fluorescent-dye conjugated secondary antibody, while brain sections labeled with GFAP or MBP were incubated with HRP secondary antibody for 4 hours. Brain sections stained with Ki67, CC3 or CD8 were washed in PBS 3 times before being mounted onto microscope slides and coverslipped with ProLong Gold. High magnification images at 63X were obtained using confocal microscopy (Carl Zeiss: MIC-System) and stains were quantified using ImageJ software. Brain sections stained with MBP and GFAP were incubated with 3, 3′-diaminobenzidine (DAB) (Biocare Medical). The reaction was quenched with 10% sodium azide; sections were washed three times in 0.1 M sodium acetate followed by dehydration in xylene. Once the slides were coverslipped with DePeX Mounting Medium (Electron Microscopy Sciences), they were imaged using brightfield setting at 20X magnification (Olympus BX53).

To quantify tumor size per mouse, vibratome brain sections from a single well were stained with Cresyl Violet as described previously (4). Approximately 5-7 tumor sections per mouse were imaged using the brightfield (Olympus BX53) setting. Area of the Nissl stain covering the tumor couture in the brightfield micrographs was quantified under the Otsu threshold using ImageJ analytical software.

Paraffin embedded 5 µm liver and brain sections were stained with H&E as described by us previously (4). Brightfield images were obtained using Olympus MA BX53 microscope.

### Complete Blood Cell Counts and Serum Biochemistry

Blood from mice was taken from the submandibular vein and transferred to EDTA coated microtainer tubes (BD Biosciences) or serum separation tubes (Biotang). For serum collection samples were left in the tubes for 20 min at room temperature to allow for blood coagulation before centrifugation at 2,000 rpm. Complete blood cell counts and serum chemistry for each sample were determined by in vivo animal core at the University of Michigan. Levels of IL-2 and IL-15 in the serum was measured by ELISA according to manufacturer’s instructions (R&D) at the Cancer Center Immunology Core, University of Michigan.

### T cell Proliferation Assay

Splenocytes from the spleens of OT-1 knockout mice (Jackson) were labeled with 5-(and 6)-carboxyfluorescein diacetate succinimidyl ester (CFSE) according the manufacturer’s instructions (ThermoFisher Scientific). Splenocytes (5.0 x 10^5^) were stimulated with 100nM SIINFEKL peptide (Anaspec) in the presence or absence of 25µM, 15mM, and 30mM D-2HG Salt (Sigma-Aldrich) for 4 days in RPMI 1640 media. T cells were stained with anti-mouse CD3 and CD8 antibodies, and proliferation was assessed by CFSE dye dilution using a flow cytometer. Levels of IFN-γ in the cultures’ supernatants was measured by ELISA according to manufacturer’s instructions (R&D) at the Cancer Center Immunology Core, University of Michigan. T cell apoptosis induced by D-2HG was identified by staining cells with Annexin V-FITC (ThermoFisher) in 1X Annexin V binding buffer and 1ug/mL propidium iodide (PI). CD8+/CD3+ T cells with positive staining for both Annexin V and PI staining were identified as dead cells. CD8+/CD3+ T cells with negative staining for both Annexin V and PI staining were identified as live cells.

### Flow cytometry

Mice were euthanized and tumors were dissected from the brain and made into single cell suspensions. Blood was subjected to Red Blood Cell lysis at room temperature for 10 minutes. Tumor infiltrating immune cells in the brain was enriched with 30%70% Percoll (GE Lifesciences) density gradient. These cells were resuspended in flow buffer and non-specific antibody binding was blocked with CD16/CD32. Dendritic cells were labeled with CD45, CD11c, CD80, CD86, MHC II and B220 antibodies. M1 macrophages were labeled with CD45 and F4/80 antibodies. M2 macrophages were labeled with CD45, F4/80 and CD206 antibodies. Tumor specific T cells were labeled with CD45, CD3, CD8 and SIINFEKL-H2Kb-tetramer. Memory T cells were labeled with CD45, CD3, CD8, CD44, and CD62L antibodies. Exhausted T cells were labeled with CD45, CD3, CD8, PD-1, TIM-3, and TIGIT antibodies. MDSCs were labeled with CD45, Gr-1 and CD11b antibodies. Tregs were labeled with CD45, CD4, CD25, and Foxp3 using the Foxp3 staining kit from BD. Intracellular IFNγ were stained using BD intracellular staining kit using the manufacturer’s instructions. Antibody staining was carried out for 30min at 4°C. Staining was performed in the following order with three washes between each step: live/dead staining, blocking, surface staining, cell fixation, intracellular staining and data acquisition. Flow data has been measured BD FACS ARIA SORP (BD Bioscience) and analyzed (Supplementary Figure 13) using Flow Jo version 10 (Treestar).

### TCGA analysis

TCGA gene expression data was analyzed using the GlioVis data visualization tool, which contains over 6500 tumor samples of approximately 50 expression datasets of a large collection of brain tumor entities (5). That includes TCGA data for WHO grade II, III and IV glioma patients. Data on IDH1 mutation status was obtained for each patient using http://gliovis. bioinfo.cnio.es portal. Cases were stratified into 2 groups on the basis of wild type (288 cases) or mutated (655 cases) IDH status. Based on genetic lesions within the tumors, they were either classified as IDH1 mutant codel (1p/19q intact), IDH1 mutant noncodel (TP53/ATRX loss), or wtIDH. Log2 gene expression levels of CD274 (PD-L1) within WT and IDH-MUT patients was compared using Tukey’s Honest Significant Difference (HSD) statistical test. For the TCGA DNA methylation analysis, two probes with enriched methylation (cg15837913 and cg19724470) within the CpG island were chosen to determine DNA methylation levels for PD-L1 based on previously described findings by *Mu et al* (6). The β-values for DNA methylation levels at the CpG sites were determined using Mexpress (7). The β-values were determined as the ratio between methylated probe signal relative to the total (methylated and unmethylated) probe signal. The β-value, ranging from 0.00 to 1.00 represents enriched methylation level of the individual CpG probe.

---

**Supplementary Figure 1:**
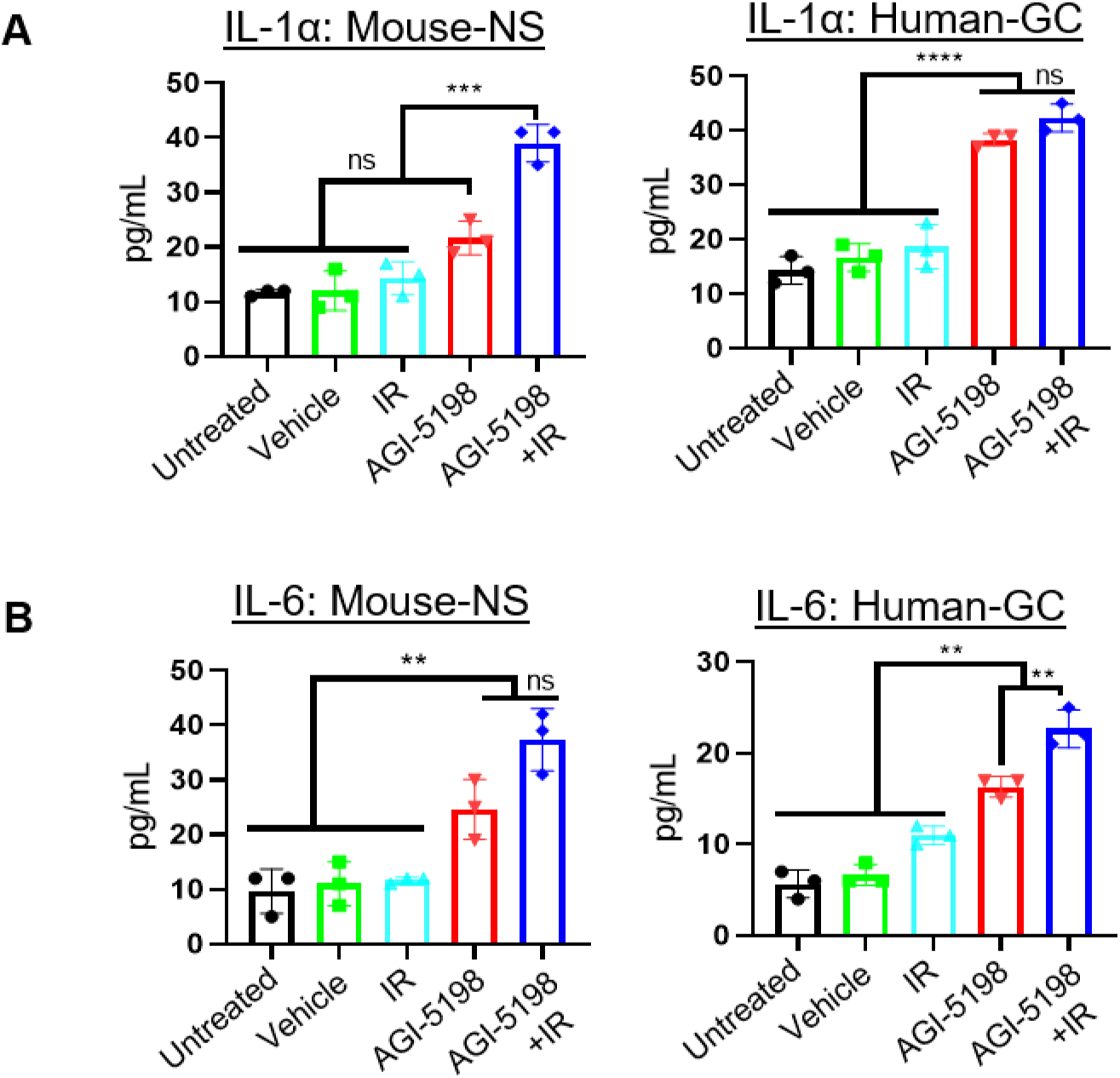
Inhibition of IDH1-R132H in combination with radiotherapy promotes the release of IL1-1α and IL-6 by mDIH1 mouse-NS and human-GC. (**A, B**) Mouse-NS were treated with 3 Gy IR in combination 1.5 μM of AGI-5198 for 72 hrs. Human-GC were treated with 10 Gy IR in combination with 5μM of AGI-5198 for 72 hrs. (A) Quantification of IL-1α release in the supernatant of mIDH1 Mouse-NS and Human-GC. (B) Quantification of IL-6 release in the supernatant of mIDH1 Mouse-NS and Human-GC. \*\**P* < 0.01, \*\*\**P* < 0.001, \*\*\*\**P* < 0.0001, one-way ANOVA test. Bars represent mean ± SEM (*n* = 3 technical replicates).

**Supplementary Figure 2:**
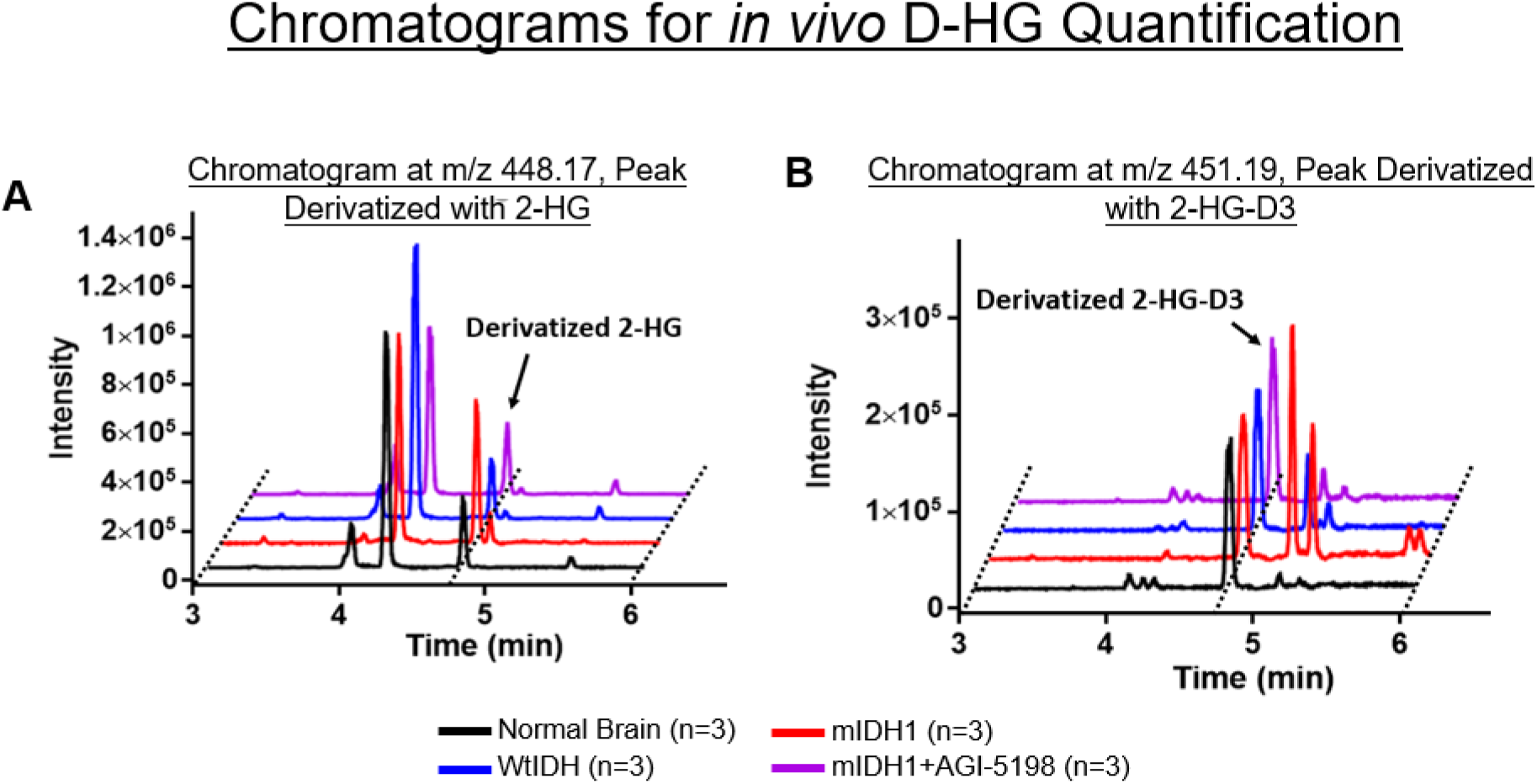
Inhibition of IDH1-R132H decreases the production of 2HG *in vivo*. UPLC-MS analysis to assess the D-2HG concentration in the TME of normal mice (blank; black), wtIDH1 glioma bearing mice (blue), mIDH1 glioma bearing mice (red), and mIDH1 glioma bearing mice treated with AGI-5198 (purple). Chromatogram to the left was acquired at m/z 448.17 (-), peak at 4.8 min is derivatized 2-HG. Chromatogram to the right was acquired at m/z 451.19 (-), peak at 4.8 min is derivatized 2-HG-D3.

**Supplementary Figure 3:**
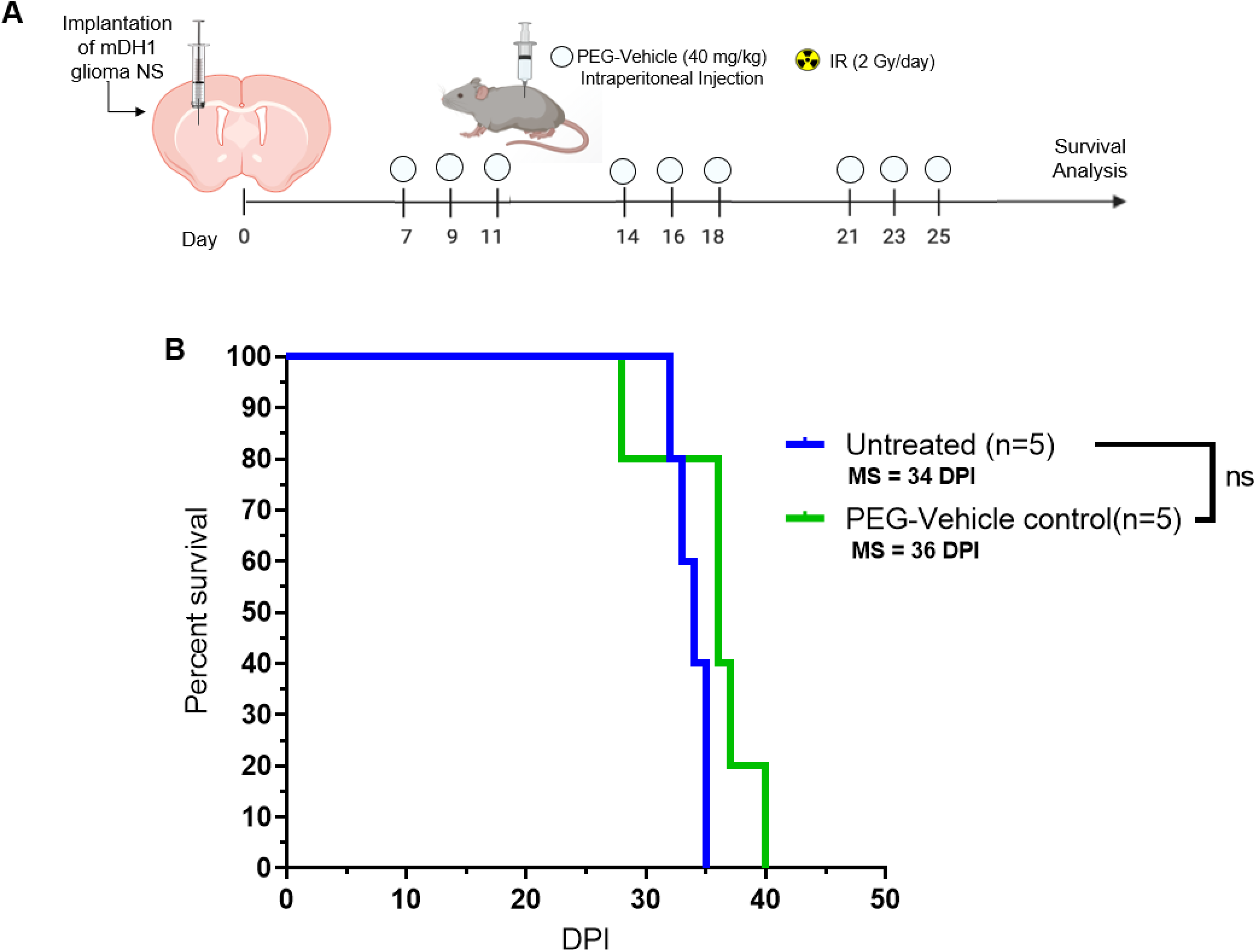
Treatment with PEG-vehicle control does not impact the survival of mIDH1 tumor bearing mice. (A) Diagram of the experimental design to assess the effect of PEG-vehicle control on the survival of mIDH1 glioma bearing mice. (**B)** Kaplan-Meier survival analysis of saline (*n* = 5) and vehicle control (*n* = 5). Data were analyzed using the log-rank (Mantel-Cox) test (ns = non-significant; MS = median survival).

**Supplementary Figure 4:**
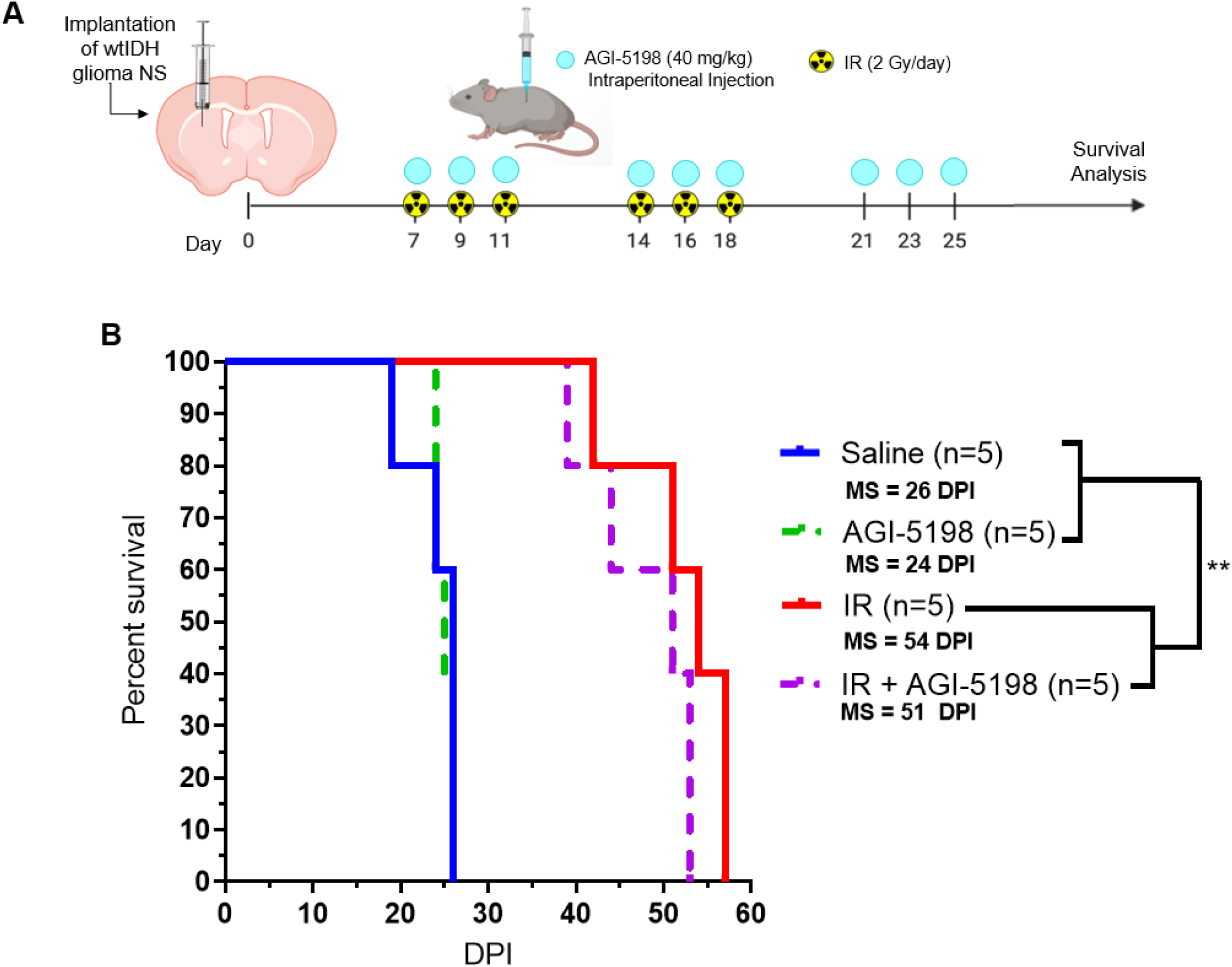
Inhibition of IDH1-R132H does not improve the survival of wtIDH1 tumor bearing mice. (A) Diagram of the experimental design to assess the impact of AGI-5198 treatment in combination with radiation on the survival of wtIDH1 glioma bearing mice. (**B)** Kaplan-Meier survival analysis of saline (*n* = 5), AGI-5198 (*n* = 5), IR (*n* = 5), and IR+AGI-5198 (*n* = 5) treated mice. Data were analyzed using the log-rank (Mantel-Cox) test. \*\**P* < 0.01; MS = median survival.

**Supplementary Figure 5:**
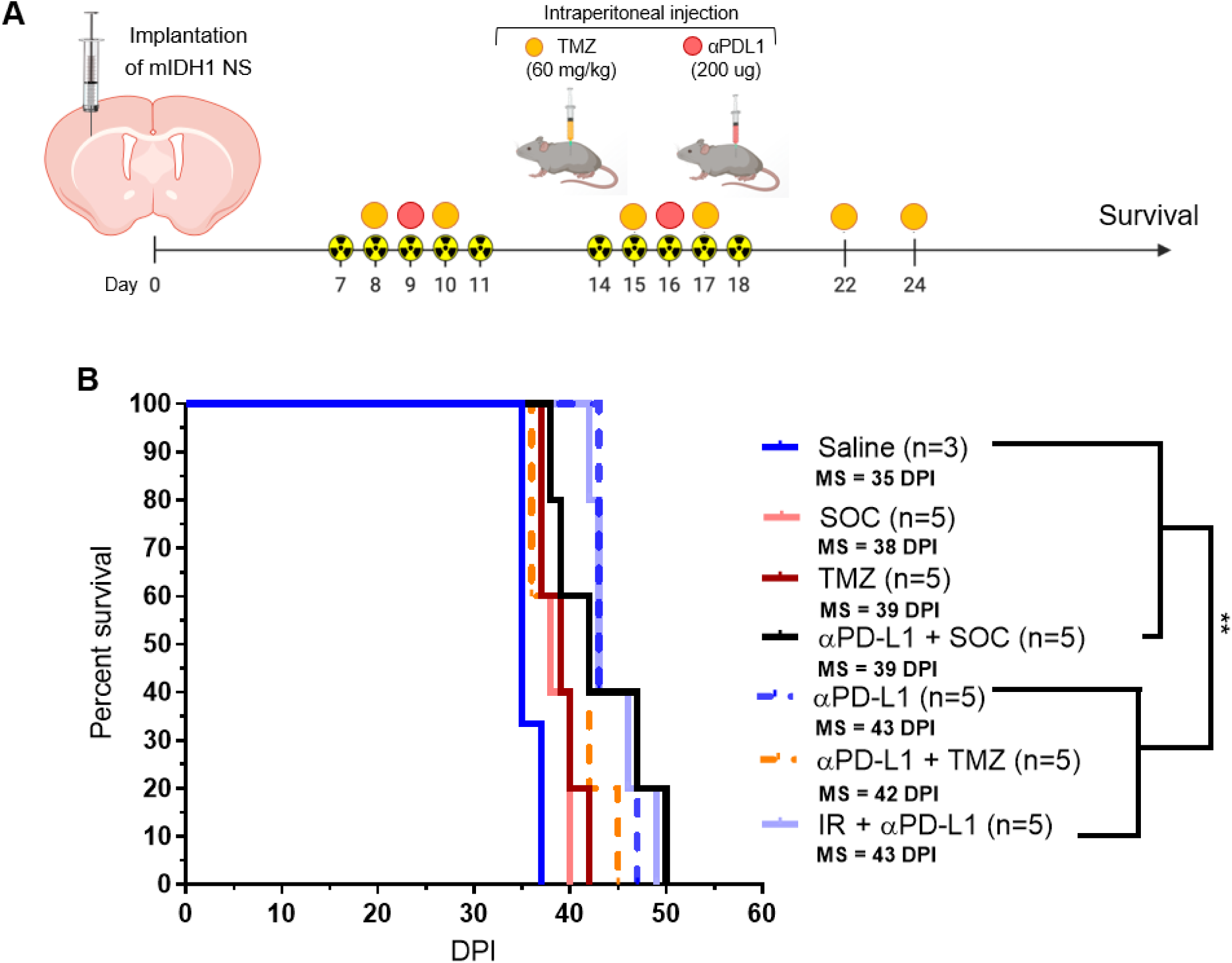
Standard of care or αPD-L1 monotherapy does not improve the survival of mIDH1 tumor bearing mice. **(A)**. Diagram of experimental design to assess the impact of SOC or αPD-L1 monotherapy on the survival of mIDH1 glioma bearing mice. (**D)** Kaplan-Meier survival analysis of saline (*n* = 3), SOC (*n* = 5), TMZ (*n* = 5), αPD-L1 + SOC (*n* = 5), αPD-L1 (*n* = 5), αPD-L1 + TMZ (*n* = 5), and IR + αPDL (*n* = 5) treated mice. Data were analyzed using the log-rank (Mantel-Cox) test. ** *P* < 0.01; MS = median survival.

**Supplementary Figure 6:**
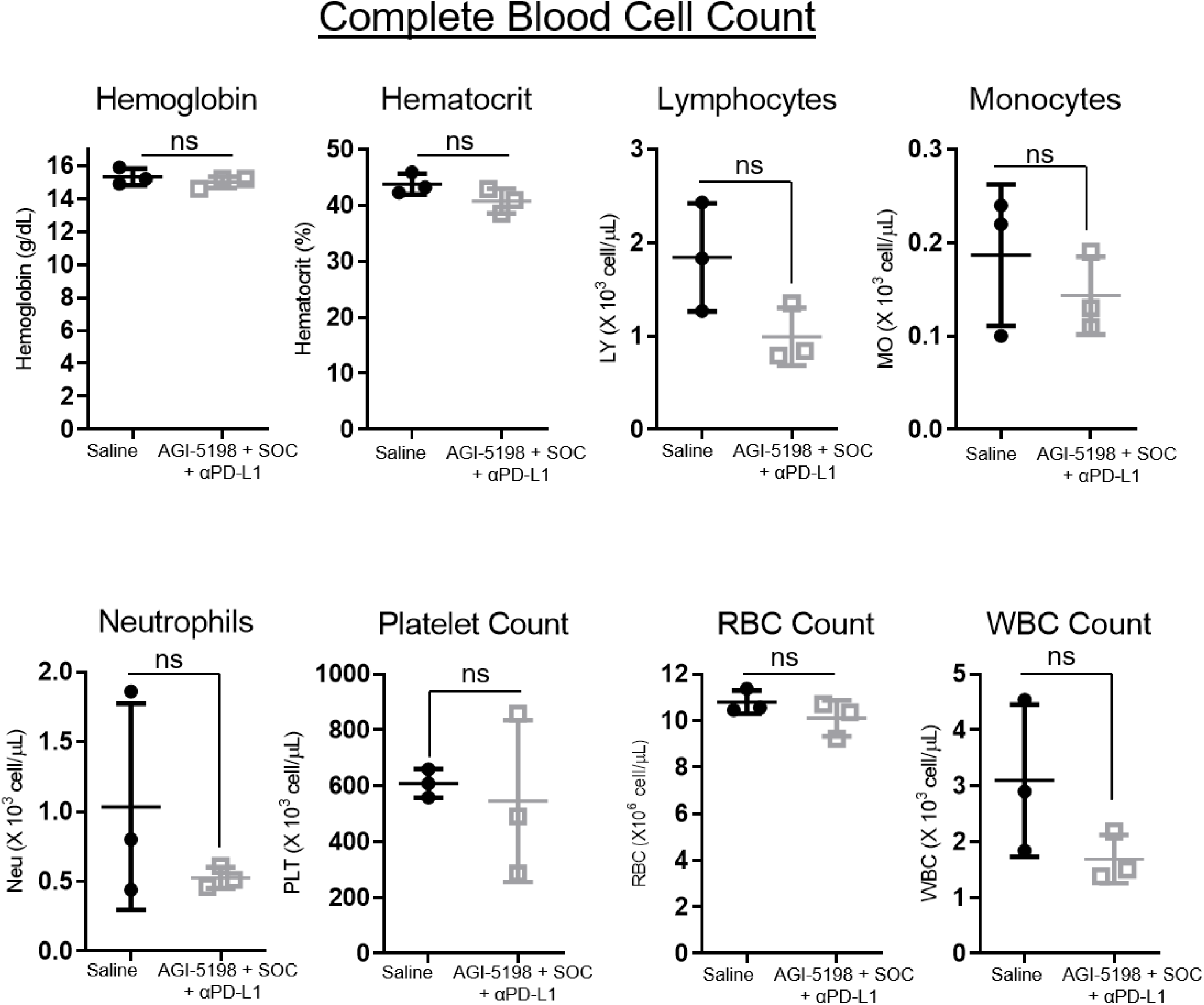
Complete blood cell (CBC) count for mIDH1 tumor bearing mice treated with IDH1-R132H inhibitor in combination with SOC and anti-PDL1 immune checkpoint blockade. Blood was collected from mIDH1 tumor bearing mice treated with saline or AGI-5198 + SOC + αPD-L1 at 27 dpi. For each treatment group levels of hemoglobin, hematocrit, lymphocytes, monocytes, neutrophils, platelet count, and red blood cell (RBC) and white blood cell (WBC) counts were quantified. The counts between saline and AGI-5198 + SOC + αPD-L1 treatment groups were compared and were non-significant, *P* > 0.05 (*n* = 3 biological replicates).

**Supplementary Figure 7:**
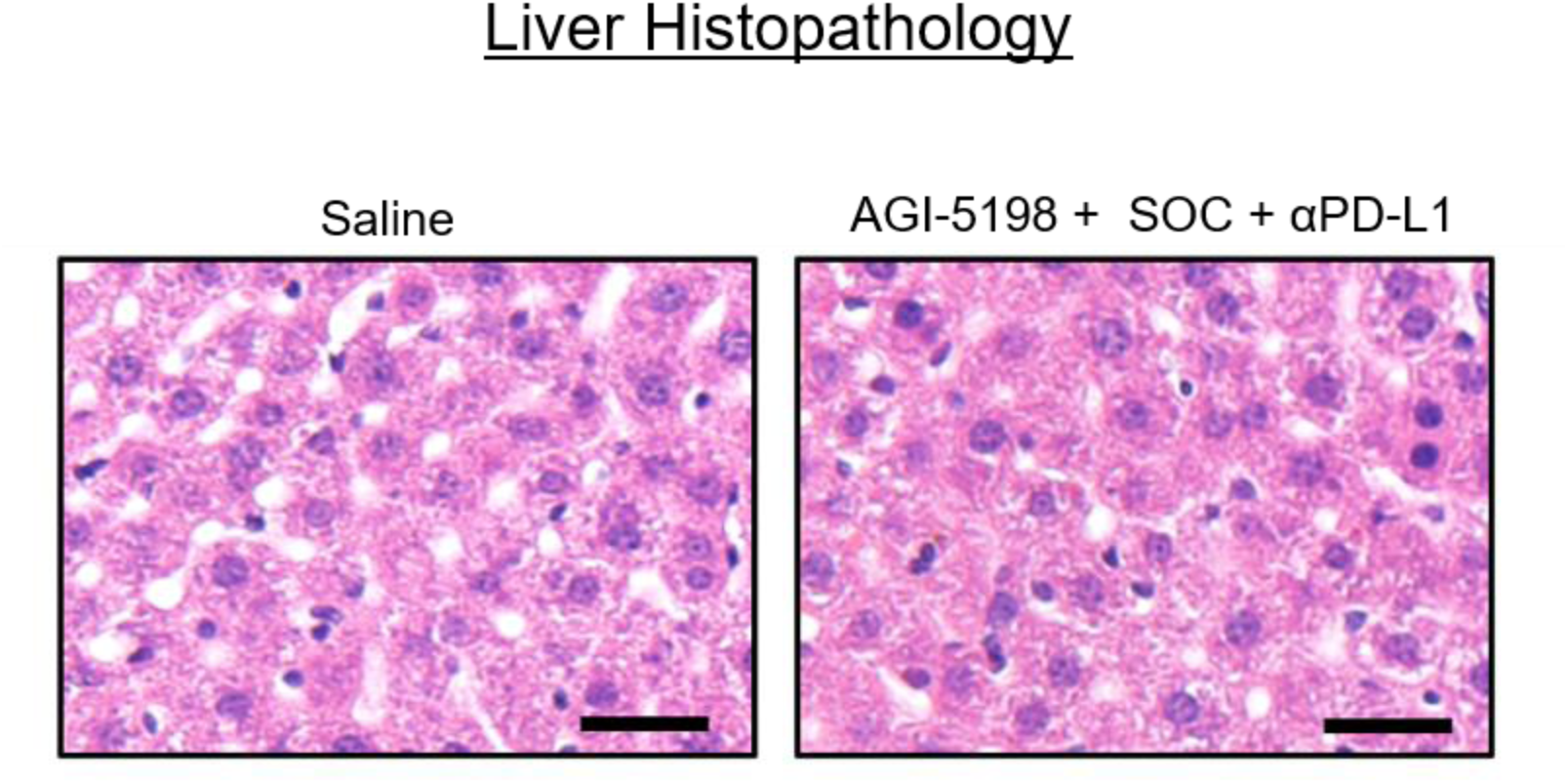
Histopathology of livers treated with IDH1-R132H inhibitor in combination with standard of care and anti-PDL1 immune checkpoint blockade. H&E staining of paraffin embedded 5 µm liver sections from mIDH1 glioma bearing mice treated with saline or AGI-5198 + SOC + αPD-L1 treatment groups at 27 dpi (representative image from *n* = 3 biological replicates, black scale bar = 100 μm).

**Supplementary Figure 8:**
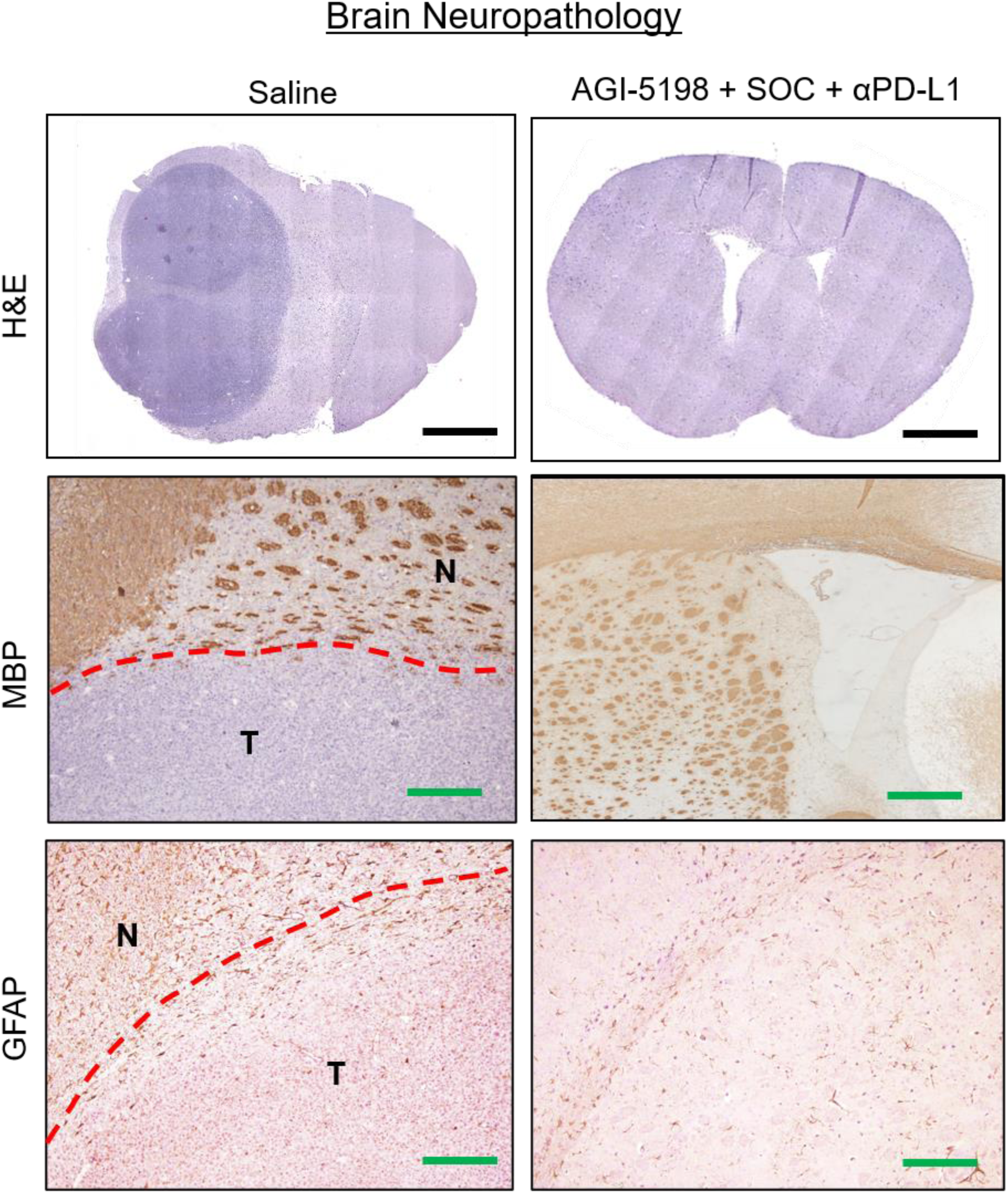
Neuropathology of brains treated with IDH1-R132H inhibitor in combination with standard of care and anti-PDL1 immune checkpoint blockade. Paraffin embedded 5 µm brain sections were obtained from saline (30 dpi), and long-term survivors in the AGI-5198 + SOC + αPD-L1 treatment groups (60 dpi after rechallenge with mIDH1 NS). Paraffin embedded 5µm brain sections from each treatment group were stained for hematoxylin and eosin (H&E), Glial fibrillary acidic protein (GFAP) or myelin basic protein (MBP). Panels show normal brain (N) and tumor (T) tissue (black scale bar = 1 mm; green scale bar = 100 μm).

**Supplementary Figure 9:**
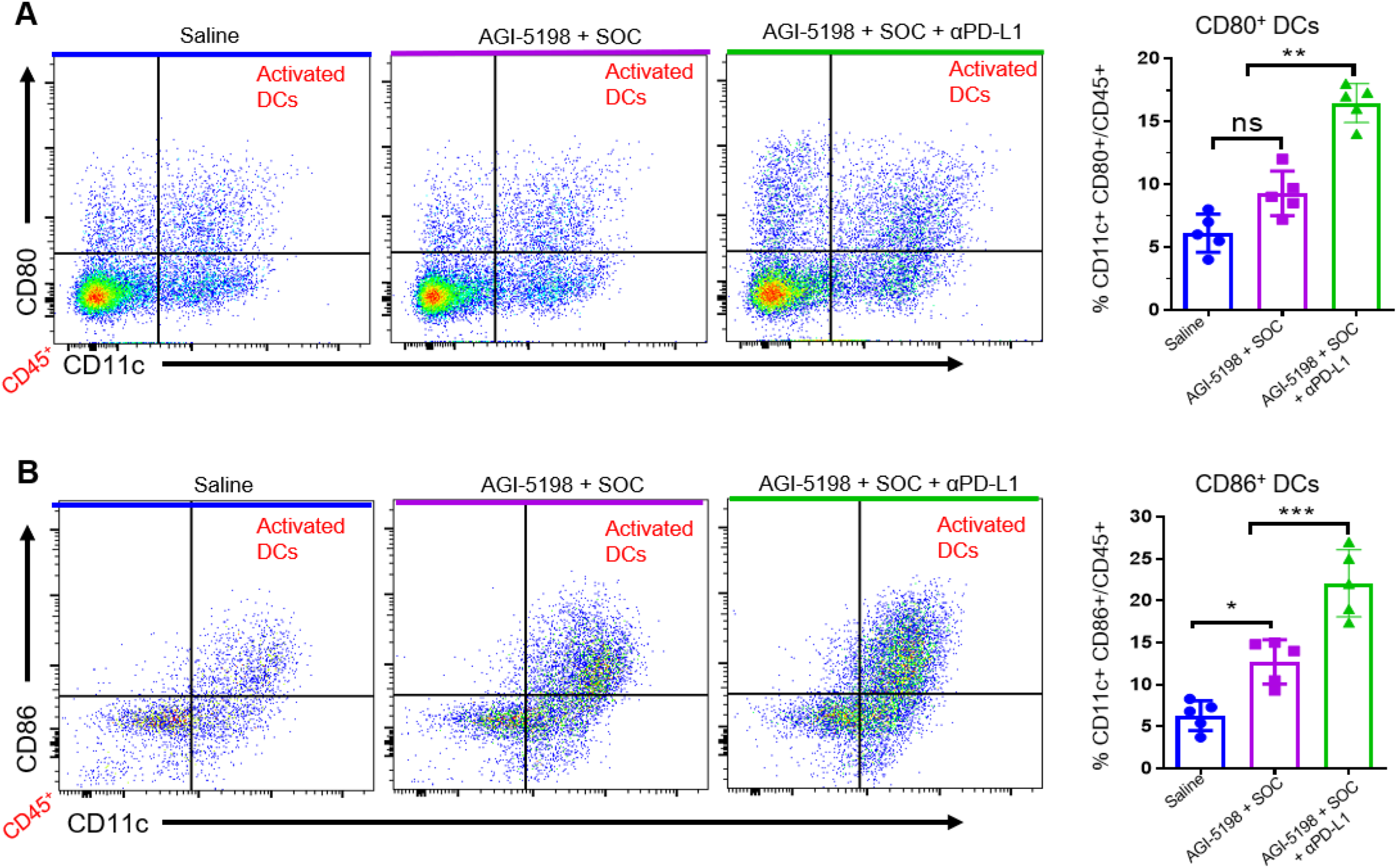
Inhibition of IDH1-R132H in combination with standard of care and anti-PDL1 blockade increases the infiltration of activated dendritic cells (DCs) in the mIDH1 glioma TME. Quantification of the percent of activated (**A**) CD80 positive and (**B**) CD86 positive DCs in the TME of saline, AGI-5198+IR, or AGI-5198 + SOC + αPD-L1 treated mIDH1 glioma bearing mice was assessed at 27 dpi. \**P* < 0.05; \*\**P* < 0.001 one-way ANOVA test. Bars represent mean ± SEM (*n* = 5 biological replicates).

**Supplementary Figure 10:**
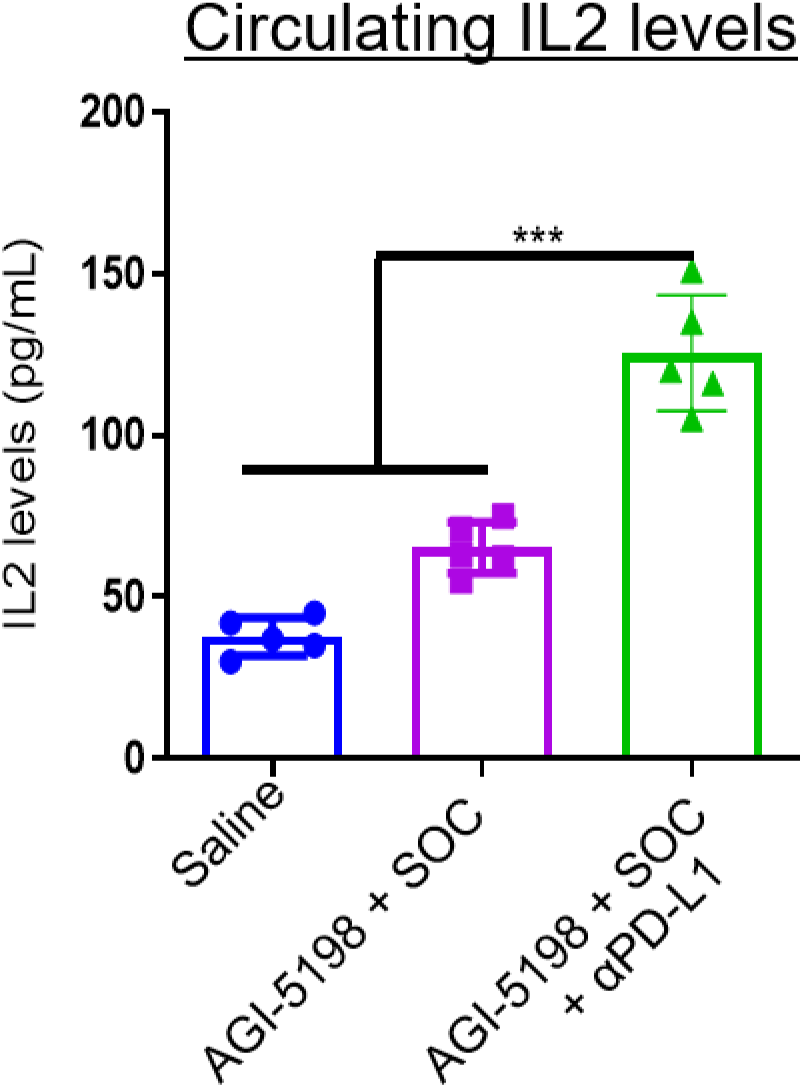
Levels of IL2 in the sera of mIDH1 tumor bearing mice treated with IDH1-R132H inhibitor in combination with standard of care and anti-PDL1 immune checkpoint blockade. (A) Quantification of IL2 in the sera of mIDH1 tumor bearing mice after treatment with saline, AGI-5198 + IR, or AGI-5198 + SOC + αPD-L1 at 27 dpi. IL2 levels were assessed by ELISA. \*\*\**P* < 0.001, one-way ANOVA test. Bars represent mean ± SEM (*n* = 5 biological replicates).

**Supplementary Figure 11:**
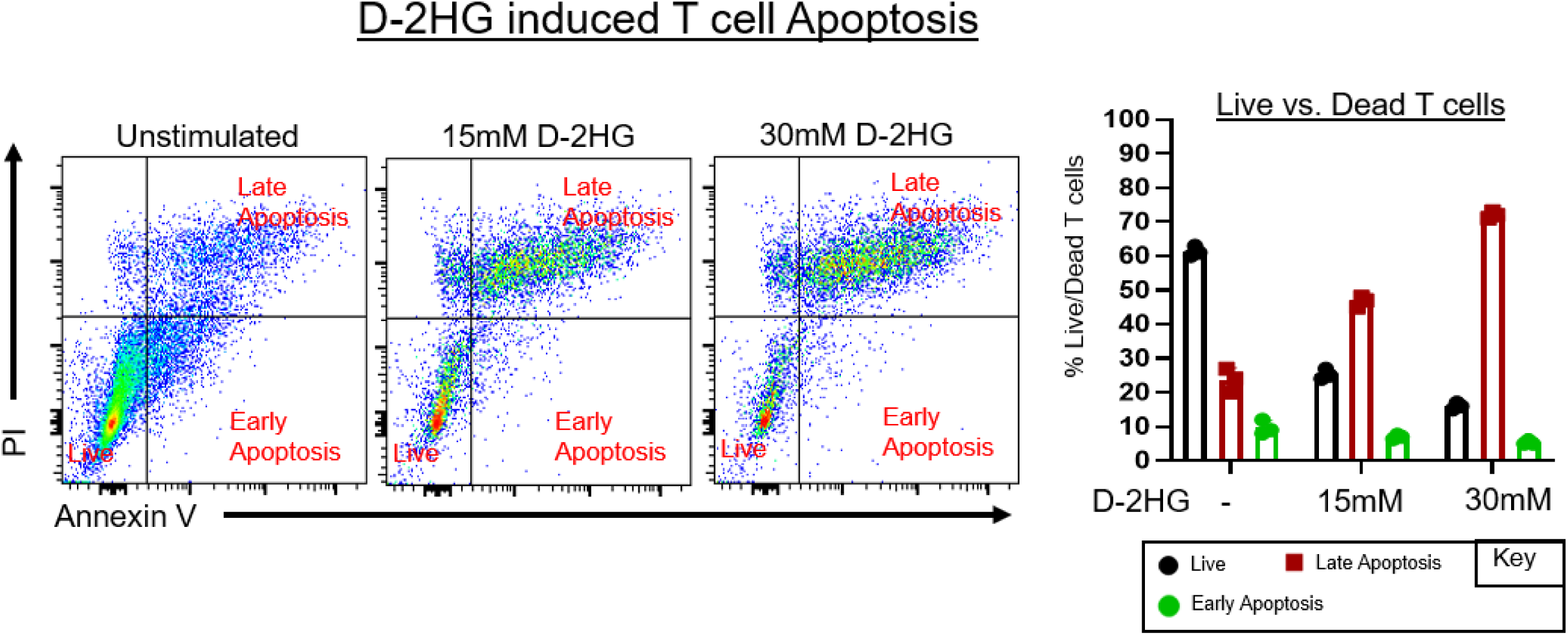
Apoptosis of T cells in response to D-2HG treatment. OT-1 splenocytes were incubated with 15mM or 30mM of D-2HG for 4 days. Then they were stained with Annexin V-FITC and propidium iodide (PI). Live non-activated T cells (CD3^+^/CD8^+^) were identified as Annexin V negative and PI negative. Dead T cells undergoing early apoptosis were identified as Annexin V positive and PI negative. Dead T cells undergoing late apoptosis were identified as Annexin V positive and PI positive.

**Supplementary Figure 12:**
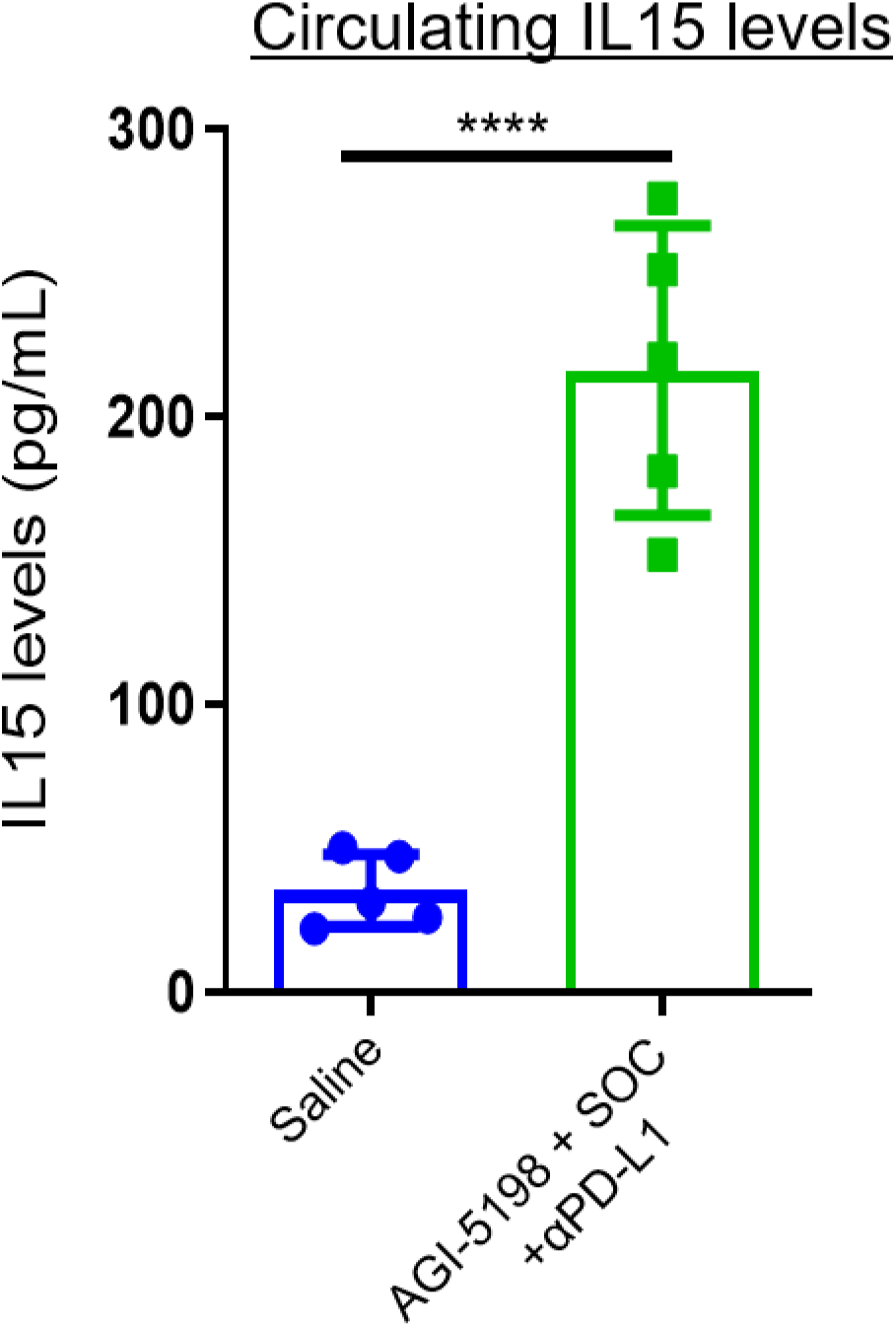
Quantification of IL15 in the sera of long-term survivors from AGI-5198 + SOC + αPD-L1 treatment rechallenged with mIDH1 glioma. Long-term survivors from the AGI-5198 + SOC + αPD-L1 treatment group were rechallenged in the contralateral hemisphere with mIDH1 NS. As a control group, untreated normal mice were implanted with mIDH1 and did not receive further treatment. Sera were collected from mice at 14 dpi after rechallenge. Quantification of IL15 in the sera of untreated or rechallenged AGI-5198 + IR + TMZ + αPD-L1 mIDH1 tumor bearing mice. IL15 levels were assessed by ELISA. \*\*\*\**P* < 0.0001, One-way ANOVA test. Bars represent mean ± SEM (*n* = 3 technical replicates).

**Supplementary Figure 13:**
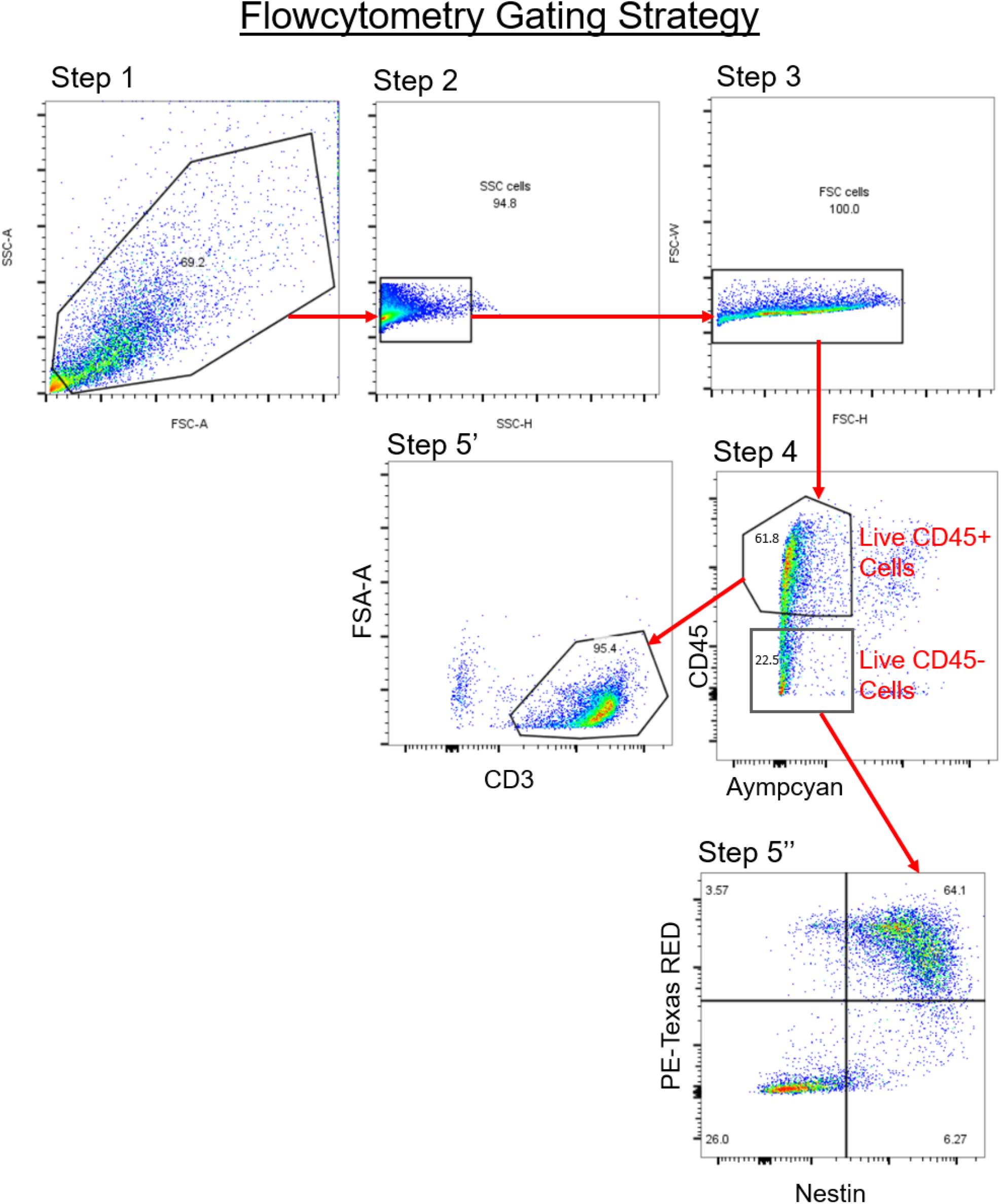
Gating strategy used for flow cytometry analysis of tumor infiltrating and peripheral lymphocytes. Step 1: Immunolabeled lymphocytes were gated to exclude cellular debris. Step 2 and Step 3: Doublet discrimination gating was performed to filter out cellular aggregates prior to analysis. Step 4: CD45+/Aymcyan-gate to identify live mIDH1 glioma infiltrating immune cells. CD45-/Aymcyan-gate to identify tumor cells. Step 5’: Identification of live CD3+ cells. Step 5: Identification of Katushka+/nestin+ live tumor cells.

## Notes

### Competing Interest Statement

The authors have declared no competing interest.

## References

1. Hadziahmetovic M, Shirai K, and Chakravarti A. Recent advancements in multimodality treatment of gliomas. Future Oncol. 2011;7(10):1169–83.

2. Ceccarelli M, Barthel FP, Malta TM, Sabedot TS, Salama SR, Murray BA, et al. Molecular Profiling Reveals Biologically Discrete Subsets and Pathways of Progression in Diffuse Glioma. Cell. 2016;164(3):550–63.

3. Louis DN, Perry A, Reifenberger G, von Deimling A, Figarella-Branger D, Cavenee WK, et al. The 2016 World Health Organization Classification of Tumors of the Central Nervous System: a summary. Acta Neuropathol. 2016;131(6):803–20.

4. Brat DJ, Verhaak RG, Aldape KD, Yung WK, Salama SR, Cooper LA, et al. Comprehensive, Integrative Genomic Analysis of Diffuse Lower-Grade Gliomas. N Engl J Med. 2015;372(26):2481–98.

5. Reiter-Brennan C, Semmler L, and Klein A. The effects of 2-hydroxyglutarate on the tumorigenesis of gliomas. Contemp Oncol (Pozn). 2018;22(4):215–22.

6. Amankulor NM, Kim Y, Arora S, Kargl J, Szulzewsky F, Hanke M, et al. Mutant IDH1 regulates the tumor-associated immune system in gliomas. Genes Dev. 2017;31(8):774–86.

7. Nunez FJ, Mendez FM, Kadiyala P, Alghamri MS, Savelieff MG, Garcia-Fabiani MB, et al. IDH1-R132H acts as a tumor suppressor in glioma via epigenetic up-regulation of the DNA damage response. Sci Transl Med. 2019;11(479).

8. Dang L, White DW, Gross S, Bennett BD, Bittinger MA, Driggers EM, et al. Cancer-associated IDH1 mutations produce 2-hydroxyglutarate. Nature. 2009;462(7274):739–44.

9. Xu W, Yang H, Liu Y, Yang Y, Wang P, Kim SH, et al. Oncometabolite 2-hydroxyglutarate is a competitive inhibitor of alpha-ketoglutarate-dependent dioxygenases. Cancer Cell. 2011;19(1):17–30.

10. Figueroa ME, Abdel-Wahab O, Lu C, Ward PS, Patel J, Shih A, et al. Leukemic IDH1 and IDH2 mutations result in a hypermethylation phenotype, disrupt TET2 function, and impair hematopoietic differentiation. Cancer Cell. 2010;18(6):553–67.

11. Chowdhury R, Yeoh KK, Tian YM, Hillringhaus L, Bagg EA, Rose NR, et al. The oncometabolite 2-hydroxyglutarate inhibits histone lysine demethylases. EMBO Rep. 2011;12(5):463–9.

12. Kaartinen V, Dudas M, Nagy A, Sridurongrit S, Lu MM, and Epstein JA. Cardiac outflow tract defects in mice lacking ALK2 in neural crest cells. Development. 2004;131(14):3481–90.

13. Turcan S, Rohle D, Goenka A, Walsh LA, Fang F, Yilmaz E, et al. IDH1 mutation is sufficient to establish the glioma hypermethylator phenotype. Nature. 2012;483(7390):479–83.

14. Urban DJ, Martinez NJ, Davis MI, Brimacombe KR, Cheff DM, Lee TD, et al. Assessing inhibitors of mutant isocitrate dehydrogenase using a suite of pre-clinical discovery assays. Sci Rep. 2017;7(1):12758.

15. Rohle D, Popovici-Muller J, Palaskas N, Turcan S, Grommes C, Campos C, et al. An inhibitor of mutant IDH1 delays growth and promotes differentiation of glioma cells. Science. 2013;340(6132):626–30.

16. Popovici-Muller J, Lemieux RM, Artin E, Saunders JO, Salituro FG, Travins J, et al. Discovery of AG-120 (Ivosidenib): A First-in-Class Mutant IDH1 Inhibitor for the Treatment of IDH1 Mutant Cancers. ACS Med Chem Lett. 2018;9(4):300–5.

17. Golub D, Iyengar N, Dogra S, Wong T, Bready D, Tang K, et al. Mutant Isocitrate Dehydrogenase Inhibitors as Targeted Cancer Therapeutics. Frontiers in Oncology. 2019;9(417).

18. Wu X, Gu Z, Chen Y, Chen B, Chen W, Weng L, et al. Application of PD-1 Blockade in Cancer Immunotherapy. Comput Struct Biotechnol J. 2019;17:661–74.

19. Kamran N, Kadiyala P, Saxena M, Candolfi M, Li Y, Moreno-Ayala MA, et al. Immunosuppressive Myeloid Cells’ Blockade in the Glioma Microenvironment Enhances the Efficacy of Immune-Stimulatory Gene Therapy. Mol Ther. 2017;25(1):232–48.

20. Altshuler DB, Kadiyala P, Nunez FJ, Nunez FM, Carney S, Alghamri MS, et al. Prospects of biological and synthetic pharmacotherapies for glioblastoma. Expert Opin Biol Ther. 2020;20(3):305–17.

21. Krysko DV, Garg AD, Kaczmarek A, Krysko O, Agostinis P, and Vandenabeele P. Immunogenic cell death and DAMPs in cancer therapy. Nat Rev Cancer. 2012;12(12):860–75.

22. Kohanbash G, Carrera DA, Shrivastav S, Ahn BJ, Jahan N, Mazor T, et al. Isocitrate dehydrogenase mutations suppress STAT1 and CD8+ T cell accumulation in gliomas. J Clin Invest. 2017;127(4):1425–37.

23. Mu L, Long Y, Yang C, Jin L, Tao H, Ge H, et al. The IDH1 Mutation-Induced Oncometabolite, 2-Hydroxyglutarate, May Affect DNA Methylation and Expression of PD-L1 in Gliomas. Front Mol Neurosci. 2018;11:82.

24. Mineharu Y, Kamran N, Lowenstein PR, and Castro MG. Blockade of mTOR signaling via rapamycin combined with immunotherapy augments antiglioma cytotoxic and memory T-cell functions. Mol Cancer Ther. 2014;13(12):3024–36.

25. Vasilevko V, Ghochikyan A, Holterman MJ, and Agadjanyan MG. CD80 (B7-1) and CD86 (B7-2) are functionally equivalent in the initiation and maintenance of CD4+ T-cell proliferation after activation with suboptimal doses of PHA. DNA Cell Biol. 2002;21(3):137–49.

26. Kitamura T, Ye G, Elliott TF, Ogra PL, Reyes VE, and Garofalo RP. Expression, Regulation and Function of the Costimulatory Molecules B7-1(CD80) and B7-2 (CD86) and MHC Class II on Human Enterocytes † 30. Pediatric Research. 1998;43(4):8-.

27. Tugues S, Burkhard SH, Ohs I, Vrohlings M, Nussbaum K, Vom Berg J, et al. New insights into IL-12-mediated tumor suppression. Cell Death Differ. 2015;22(2):237–46.

28. Quail DF, and Joyce JA. The Microenvironmental Landscape of Brain Tumors. Cancer Cell. 2017;31(3):326–41.

29. Pilipow K, Roberto A, Roederer M, Waldmann TA, Mavilio D, and Lugli E. IL15 and T-cell Stemness in T-cell-Based Cancer Immunotherapy. Cancer Res. 2015;75(24):5187–93.

30. Lu X, Liu J, Cui P, Liu T, Piao C, Xu X, et al. Co-inhibition of TIGIT, PD1, and Tim3 reverses dysfunction of Wilms tumor protein-1 (WT1)-specific CD8+ T lymphocytes after dendritic cell vaccination in gastric cancer. Am J Cancer Res. 2018;8(8):1564–75.

31. Chaturvedi A, Herbst L, Pusch S, Klett L, Goparaju R, Stichel D, et al. Pan-mutant-IDH1 inhibitor BAY1436032 is highly effective against human IDH1 mutant acute myeloid leukemia in vivo. Leukemia. 2017;31(10):2020–8.

32. Tateishi K, Wakimoto H, Iafrate AJ, Tanaka S, Loebel F, Lelic N, et al. Extreme Vulnerability of IDH1 Mutant Cancers to NAD+ Depletion. Cancer Cell. 2015;28(6):773–84.

33. Garrett M, Sperry J, Braas D, Yan W, Le TM, Mottahedeh J, et al. Metabolic characterization of isocitrate dehydrogenase (IDH) mutant and IDH wildtype gliomaspheres uncovers cell type-specific vulnerabilities. Cancer Metab. 2018;6:4.

34. Subramanian A, Tamayo P, Mootha VK, Mukherjee S, Ebert BL, Gillette MA, et al. Gene set enrichment analysis: a knowledge-based approach for interpreting genome-wide expression profiles. Proc Natl Acad Sci U S A. 2005;102(43):15545–50.

35. Koschmann C, Calinescu AA, Nunez FJ, Mackay A, Fazal-Salom J, Thomas D, et al. ATRX loss promotes tumor growth and impairs nonhomologous end joining DNA repair in glioma. Sci Transl Med. 2016;8(328):328ra28.

36. Herceg Z. Epigenetic Mechanisms as an Interface Between the Environment and Genome. Adv Exp Med Biol. 2016;903:3–15.

37. Koschmann C, Nunez FJ, Mendez F, Brosnan-Cashman JA, Meeker AK, Lowenstein PR, et al. Mutated Chromatin Regulatory Factors as Tumor Drivers in Cancer. Cancer Res. 2017;77(2):227–33.

38. Leu S, von Felten S, Frank S, Boulay JL, and Mariani L. IDH mutation is associated with higher risk of malignant transformation in low-grade glioma. J Neurooncol. 2016;127(2):363–72.

39. Parsons DW, Jones S, Zhang X, Lin JC, Leary RJ, Angenendt P, et al. An integrated genomic analysis of human glioblastoma multiforme. Science. 2008;321(5897):1807–12.

40. Kamran N, Alghamri MS, Nunez FJ, Shah D, Asad AS, Candolfi M, et al. Current state and future prospects of immunotherapy for glioma. Immunotherapy. 2018;10(4):317–39.

41. Weenink B, Draaisma K, Ooi HZ, Kros JM, Sillevis Smitt PAE, Debets R, et al. Low-grade glioma harbors few CD8 T cells, which is accompanied by decreased expression of chemo-attractants, not immunogenic antigens. Sci Rep. 2019;9(1):14643.

42. Bunse L, Pusch S, Bunse T, Sahm F, Sanghvi K, Friedrich M, et al. Suppression of antitumor T cell immunity by the oncometabolite (R)-2-hydroxyglutarate. Nat Med. 2018;24(8):1192–203.

43. Fu X, Chin RM, Vergnes L, Hwang H, Deng G, Xing Y, et al. 2-Hydroxyglutarate Inhibits ATP Synthase and mTOR Signaling. Cell Metab. 2015;22(3):508–15.

44. Bottcher M, Renner K, Berger R, Mentz K, Thomas S, Cardenas-Conejo ZE, et al. D-2-hydroxyglutarate interferes with HIF-1alpha stability skewing T-cell metabolism towards oxidative phosphorylation and impairing Th17 polarization. Oncoimmunology. 2018;7(7):e1445454.

45. Ye D, Guan KL, and Xiong Y. Metabolism, Activity, and Targeting of D- and L-2- Hydroxyglutarates. Trends Cancer. 2018;4(2):151–65.

46. Kroemer G, Galluzzi L, Kepp O, and Zitvogel L. Immunogenic cell death in cancer therapy. Annu Rev Immunol. 2013;31:51–72.

47. Hou W, Zhang Q, Yan Z, Chen R, Zeh Iii HJ, Kang R, et al. Strange attractors: DAMPs and autophagy link tumor cell death and immunity. Cell Death Dis. 2013;4:e966.

48. Ko A, Kanehisa A, Martins I, Senovilla L, Chargari C, Dugue D, et al. Autophagy inhibition radiosensitizes in vitro, yet reduces radioresponses in vivo due to deficient immunogenic signalling. Cell Death Differ. 2014;21(1):92–9.

49. Koukourakis MI, Mitrakas AG, and Giatromanolaki A. Therapeutic interactions of autophagy with radiation and temozolomide in glioblastoma: evidence and issues to resolve. Br J Cancer. 2016;114(5):485–96.

50. Kim ES, Kim JE, Patel MA, Mangraviti A, Ruzevick J, and Lim M. Immune Checkpoint Modulators: An Emerging Antiglioma Armamentarium. J Immunol Res. 2016;2016:4683607.

## References

1. Tateishi K, Wakimoto H, Iafrate AJ, Tanaka S, Loebel F, Lelic N, et al. Extreme Vulnerability of IDH1 Mutant Cancers to NAD+ Depletion. Cancer Cell. 2015;28(6):773–84.

2. Nunez FJ, Mendez FM, Kadiyala P, Alghamri MS, Savelieff MG, Garcia-Fabiani MB, et al. IDH1-R132H acts as a tumor suppressor in glioma via epigenetic up-regulation of the DNA damage response. Sci Transl Med. 2019;11(479).

3. Leder K, Pitter K, LaPlant Q, Hambardzumyan D, Ross BD, Chan TA, et al. Mathematical modeling of PDGF-driven glioblastoma reveals optimized radiation dosing schedules. Cell. 2014;156(3):603–16.

4. Calinescu AA, Yadav VN, Carballo E, Kadiyala P, Tran D, Zamler DB, et al. Survival and Proliferation of Neural Progenitor-Derived Glioblastomas Under Hypoxic Stress is Controlled by a CXCL12/CXCR4 Autocrine-Positive Feedback Mechanism. Clin Cancer Res. 2017;23(5):1250–62.

5. Bowman RL, Wang Q, Carro A, Verhaak RG, and Squatrito M. GlioVis data portal for visualization and analysis of brain tumor expression datasets. Neuro Oncol. 2017;19(1):139–41.

6. Mu L, Long Y, Yang C, Jin L, Tao H, Ge H, et al. The IDH1 Mutation-Induced Oncometabolite, 2-Hydroxyglutarate, May Affect DNA Methylation and Expression of PD-L1 in Gliomas. Front Mol Neurosci. 2018;11:82.

7. Wei J, Li G, Zhang J, Zhou Y, Dang S, Chen H, et al. Integrated analysis of genome-wide DNA methylation and gene expression profiles identifies potential novel biomarkers of rectal cancer. Oncotarget. 2016;7(38):62547–58.

8. Garcia-Fabiani M, Kadiyala P, Lowenstein PR, Castro MG. An optimized portocol for in vivo analysis of tumor cell division in a Sleeping Beauty-mediated mouse glioma model. STAR Protocols. 2020; In Press.

